# Engineering quantitative root disease resistance in barley by targeting conserved SCAR susceptibility genes without compromising seed yield or mycorrhizal symbiosis

**DOI:** 10.64898/2026.03.31.715550

**Authors:** Sabine Brumm, Matthew Macleod, Isaac Coven, Inmaculada Hernándes-Pinzón, Edouard Evangelisti, Marion C. Müller, Matthew J. Moscou, Sebastian Schornack

## Abstract

Durable resistance to soil-borne pathogens remains elusive in cereals, partly because susceptibility (S) genes that facilitate root infection have not been identified in monocots. In the model legume *Medicago truncatula*, the SCAR/WAVE complex member *MtAPI* functions as a root S-gene for microbial invasion. Whether *SCAR* gene associated susceptibility function is conserved in monocots, and whether *SCAR* gene inactivation can enhance root resistance in cereals, remains unknown. Here, we identify and characterize three SCAR genes in barley: *HvSCAR-A*, *HvSCAR-B*, and *HvSCAR-C*. Cross-species complementation assays indicate that *HvSCAR-B* and *HvSCAR-C* are functionally similar to *MtAPI*. While *hscar-b* and *hvscar-c* single mutants exhibited no major growth defects, *hvscar-a* mutants showed strongly reduced seed production, and a *hvscar-b/c* double mutant displayed shorter root hairs. Notably, the *hvscar-b/c* double mutant exhibited increased resistance to the hemibiotrophic pathogen *Phytophthora palmivora* but greater colonization by the symbiotic arbuscular mycorrhizal fungus *Funneliformis mosseae*, underscoring a complex role in plant root - microbe interactions. Our findings reveal a conserved susceptibility function of SCAR genes in monocots and identify *api* monocot homologs as promising targets for engineering disease resistance in cereals. This study offers new insights into SCAR protein functional diversification and its potential for improving root health in crop plants.

## Introduction

Plant disease management remains a critical challenge in agriculture and traditionally relies on resistance (*R*) genes and chemical treatments (Fisher et al., 2018; Jones et al., 2024). However, *R* gene-mediated resistance is often pathogen-specific and can be rapidly overcome by evolving pathogens (McDonald and Linde, 2002). Chemical control can also be compromised by the development of resistance in pathogen populations (Fisher et al., 2018) and is generally most effective against foliar diseases, whereas soil-borne pathogens remain difficult to control. With climate change expected to exacerbate the prevalence and severity of soil-borne diseases (Delgado-Baquerizo et al., 2020), there is an urgent need for alternative disease management strategies. Targeted removal or modification of susceptibility (*S*) genes, host plant genes that facilitate microbial compatibility, offers a promising alternative for durable resistance (van Schie and Takken, 2014; Garcia-Ruiz et al., 2021) However, *S* genes can also contribute to mutualistic interactions (Rey et al., 2015) and plant development (Jørgensen, 1992; Pathuri et al., 2008; Scheler et al., 2016), therefore it is critical to identify those whose loss does not compromise key agronomic traits such as growth and yield.

Barley (*Hordeum vulgare*, *Hv*), the fourth most produced cereal crop worldwide (FAOSTAT, 2024), is cultivated for diverse uses including malt production, baking, and animal feed (Newton et al., 2011). As a hardy grain, it can be grown in environments too harsh for many other cereals (Abdelghany et al., 2024). One of the most successful examples of durable disease resistance deployment via S-gene modification in barley are the recessive *mlo* alleles, which have been deployed globally to confer near-complete resistance to the foliar powdery mildew disease caused by *Blumeria hordei* (Freisleben and Lein, 1942; Büschges et al., 1997; Jørgensen and Wolfe, 2011). Knockout of *MLO* is accompanied by pleiotropic phenotypes such as spontaneous leaf lesioning (Jørgensen, 1992) and increased susceptibility to other pathogens (Jarosch et al., 1999), yet its agronomic benefits have supported long-term, widespread use (Jørgensen, 1992). In contrast, comparable *S*-gene-based strategies for the control of economically relevant root-rot diseases caused by fungi and oomycetes (Gutteridge et al., 2003; Schroeder and Paulitz, 2006; Basheer et al., 2022; Liu et al., 2024) are currently lacking, largely because no susceptibility genes have yet been identified or validated in barley roots.

Beyond pathogenic interactions, barley roots establish endosymbiotic associations with arbuscular mycorrhizal fungi (AMF), marked by the intracellular growth of fungal hyphae and the formation of specialized structures such as cortical arbuscules and lipid storage vesicles (Choi et al., 2018). AMF symbiosis supports a bidirectional plant-fungus exchange in which the plant supplies carbohydrates and lipids to the fungus in return for mineral nutrients and water delivered via extensive extraradical hyphal networks that extend root foraging capacity beyond the rhizosphere (Jakobsen and Rosendahl, 1990; Keymer et al., 2017; Luginbuehl et al., 2017; Sawers et al., 2017; Kakouridis et al., 2022). Because AMF generally enhance plant nutrient acquisition, improve stress tolerance (Chitarra et al., 2016; Berger and Gutjahr, 2021), and contribute to protection against soil-borne diseases (Campo et al., 2020; Guyon et al., 2025), they are increasingly recognized as valuable biofertilizers in sustainable agriculture. It is therefore critical to assess whether S-gene manipulation affects AMF colonization. Although barley’s mycorrhizal receptivity has generally been reported as low (Grace et al., 2009), recent experimental data show that AMF can significantly enhance nutrient acquisition under both ambient and elevated CO₂ conditions in barley (Thirkell et al., 2021). Notably, cereals and *Medicago truncatula* (*M. truncatula*, *Mt*) carrying *mlo* mutations exhibit reduced early AMF colonization and lower induction of symbiosis-related genes (Jacott et al., 2020), highlighting that altering susceptibility genes can compromise AM symbiosis. This raises the broader question of whether other susceptibility genes can be modified to enhance pathogen resistance without introducing similar trade-offs.

Previously, the *SCAR* gene *MtAPI* was identified as a root susceptibility factor in the model legume *M. truncatula*, where it modulates interactions with both, beneficial nitrogen-fixing rhizobia and the pathogenic oomycete *Phytophthora palmivora* (Teillet et al., 2008; Gavrin et al., 2020). *P. palmivora* represents an aggressive hemibiotrophic pathogen responsible for severe diseases in numerous tropical and subtropical crops. It exhibits a broad host range, colonizing both root and foliar tissues across diverse plant lineages, including economically relevant dicots such as tomato (*Solanum lycopersicum*) (Fawke et al., 2019) and monocots such as barley (Le Fevre et al., 2016). In *M. truncatula*, the contribution of *MtAPI* to host susceptibility is linked to its role within the conserved SCAR/WAVE complex, where it influences actin-dependent polarized secretion. Loss of *MtAPI* function perturbs actin remodeling and selectively impairs the deposition of cell wall polysaccharides, including xyloglucan, rhamnogalacturonan I, and homogalacturonan. These alterations in wall architecture quantitatively increase root resistance to *P. palmivora* infection (Gavrin et al., 2020) but do not impact AMF colonisation in *M. truncatula api* roots (Teillet et al., 2008). Whether this development-linked resistance mechanism is conserved beyond dicots and can be applied in monocots remains unclear.

Despite exhibiting shorter root hairs, *api* mutants retain normal growth and reproduction (Gavrin et al., 2020), making *SCAR* genes attractive candidates for *S*-gene-based resistance engineering. The SCAR/WAVE complex is a highly conserved regulator of actin dynamics that promotes the assembly of branched actin networks by activating the ARP2/3 complex (Yanagisawa et al., 2013). Plant SCAR/WAVE proteins carry an N-terminal SCAR-homology domain (SHD) required for complex assembly and a C-terminal Wiskott–Aldrich homology 2, central, and acidic domain (WCA) that stimulates ARP2/3-dependent actin nucleation (Yanagisawa et al., 2013). Mutations in SCAR/WAVE components frequently disrupt cell morphology, anisotropic cell expansion, and cell–cell adhesion and contribute to specialized developmental processes, including trichome formation, leaf-shape regulation, and root-hair elongation (Li et al., 2004; Basu et al., 2005; Djakovic et al., 2006; Uhrig et al., 2007; Miyahara et al., 2010). Notably, SCARs are the only members of the SCAR/WAVE complex that have undergone expansion into multigene families in plants, enabling paralogs to diversify and acquire distinct, tissue-specific roles. Such functional diversification is illustrated in *M. truncatula*, where *MtAPI* and its closely related paralog *MtHAPI1* fulfill non-redundant functions (Brumm et al., 2025). In monocots, however, *SCAR* gene function remains poorly characterized: the only genetically defined example to date is *TUT1* in rice which is essential for panicle development and seed yield (Bai et al., 2015). Whether SCAR homologs in monocots are all specialized for reproductive development or have diversified in function is not known yet. This gap in knowledge raises the question of whether targeting SCAR homologs in cereals can confer resistance to root-infecting pathogens without unwanted effects on growth and yield.

In this study, we identify and functionally characterise three previously unstudied barley *SCAR* gene family members and show that their proteins fall into two major angiosperm SCAR clades. Our cross-species mutant complementation experiments suggest conserved functionality among clade II SCAR genes. CRISPR-Cas9 single frame-shift or deletion mutants in barley further support clade-specific roles: clade II *hvscar*-*b* and *hvscar*-*c* plants show no major vegetative defects, clade I *hvscar*-*a* mutants display reduced seed production. Under the tested conditions *hvscar*-*b,c* double mutant plants exhibited enhanced root resistance to *P. palmivora*. In contrast to their conserved role in promoting pathogen susceptibility, loss of *HvSCAR-B* and *HvSCAR-C* led to increased arbuscular mycorrhiza colonisation, indicating that clade II HvSCAR genes affect pathogenic and beneficial interactions differently. Taken together, we demonstrate that *HvSCAR* genes perform clade-specific roles in development and root-microbe interactions, respectively. Our results highlight clade II SCAR genes as promising candidates for future S-gene engineering in cereals.

## Results

### *HvSCAR*-*B* and *HvSCAR*-*C*, but not *HvSCAR*-*A*, functionally complement the *M. truncatula api* root-hair defect

To test whether SCAR-mediated susceptibility mechanisms described for *M. truncatula* extend to cereals, we identified SCAR homologs in barley and subsequently tested which, if any, retain *Mt*API-like function. Using the *Mt*API protein as a reference, we searched publicly available plant proteomes for candidate proteins that (i) carried an N-terminal SHD, and (ii) contained a C-terminal WCA. Applying these criteria, we curated a set of 429 orthologous SCAR sequences from 135 species across 74 plant families (Data S1). The analysis recovered SCAR homologs across land plants and supports an early duplication within angiosperms that resolved into two major clades (Figure S1). The first clade (CI), comprises *M. truncatula Mt*HAPI2, *Arabidopsis thaliana (A. thaliana; At) At*SCAR1/*At*SCAR3/*At*WAVE5, and a set of monocot sequences we term monocot subclade A, including barley *Hv*SCAR-A. The second clade (CII), contains lineage-specific expansions in *Fabaceae*, *Brassicaceae*, and monocots: *Fabaceae* split into *Mt*API-like and *Mt*HAPI1-like subclades; *Brassicaceae* into *A. thaliana At*SCAR2- and *At*SCAR4-related groups; and monocots into two subclades with barley *Hv*SCAR-B and *Hv*SCAR-C, respectively. The N-terminal SHD and C-terminal WCA domains in all three *Hv*SCARs are highly conserved, whereas the central regions show high divergence (Figures S2 and S3a), consistent with patterns reported for *A. thaliana* and *M. truncatula* SCARs (Kollmar et al., 2012; Brumm et al., 2025). Mining the PANBARLEX barley pangenome expression atlas, we found that all three *HvSCAR* genes are expressed across embryo, root, shoot, inflorescence, and caryopsis tissues in cv. Golden Promise, with *HvSCAR*-*B* generally exhibiting the lowest transcript abundance (Figure S3b).

**Figure 1:**
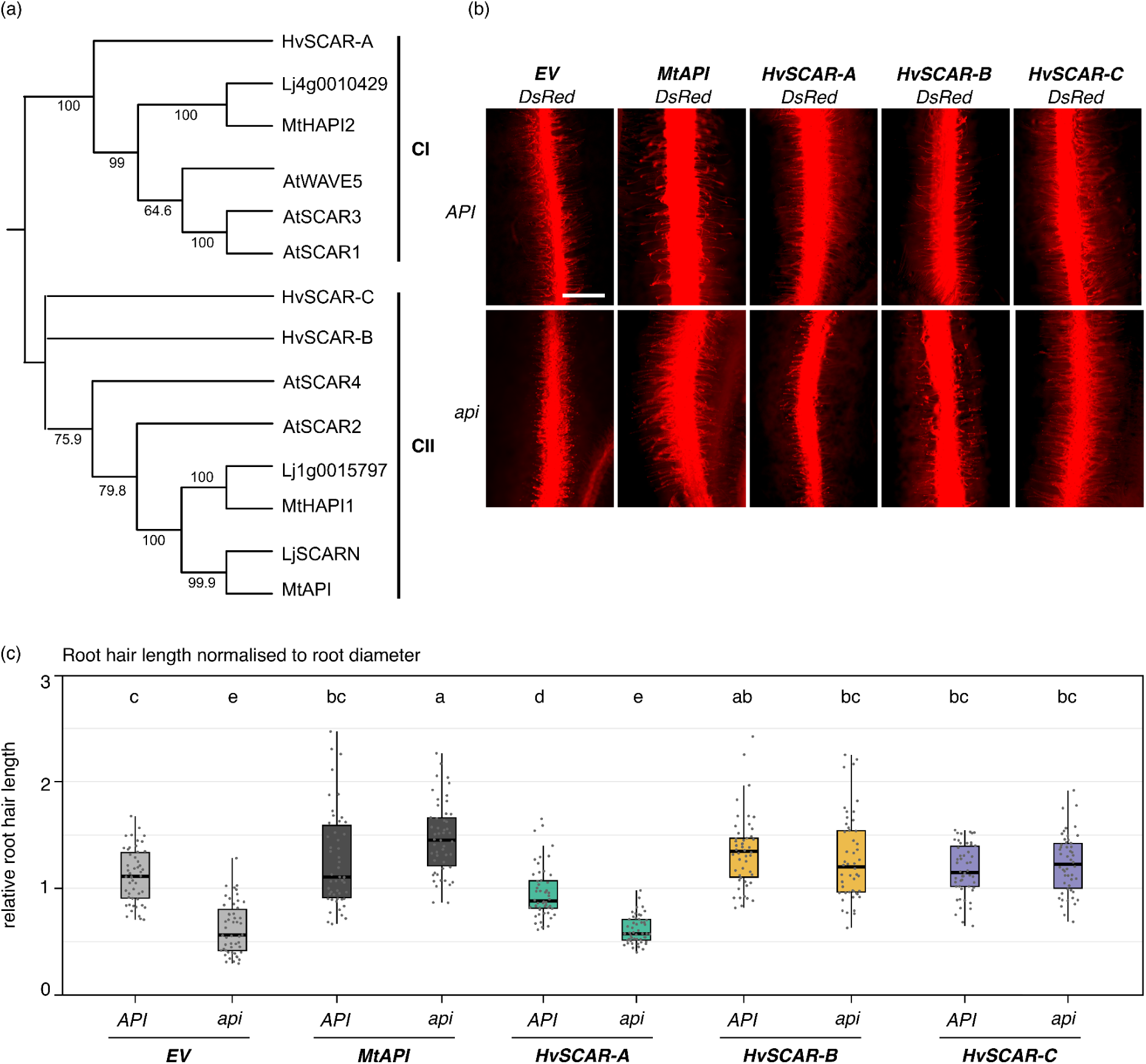
HvSCARs exhibit clade-linked functionality in *api* root-hair complementations. (a) Minimum phylogenetic tree with *Hordeum vulgare (Hv)*, *Medicago truncatula (Mt), Lotus japonicus (Lj)* and *Arabidopsis thaliana (At)* SCAR proteins. Clade I (CI) and clade II (CII) are marked by bold black lines. (b) Epifluorescence microscopy images of *M. truncatula* A17 (wild type) and *api* mutant hairy roots expressing DsRed only (pAtUBQ10:DsRed, empty vector control = EV), or together with *MtAPI*, *HvSCAR-A*, *HvSCAR-B*, or *HvSCAR-C,* all driven by the *MtAPI* promoter. All images were acquired with the same magnification, scale bar: 0.5 mm. (c) Boxplot of normalized root hair length. The root hair length (in mm) was normalized to root diameter (in mm). Each dot represents a single root hair/root diameter ratio (n = 50 per genotype). For each genotype, 10 root hairs were measured from 5 independently transformed roots. Statistical analysis: Shapiro-Wilk test, followed by Kruskal-Wallis test with Bonferroni correction. Statistical significance groupings are indicated by letters.

To test whether any of the identified *Hv*SCAR proteins can functionally replace *Mt*API, we used the root-hair elongation defect of the *M. truncatula api* mutant as a rapid, scorable readout. Prior work showed that the short-root-hair phenotype of *api* can be complemented by other SCAR CII-clade members (Figure 1a), including *A. thaliana At*SCAR2 and *Lotus japonicus (Lj)* homolog *Lj*SCARN (Gavrin et al., 2020), establishing root-hair elongation as a sensitive proxy for *Mt*API-like function. Using a 2-kb *MtAPI* promoter fragment (Gavrin et al., 2020), we expressed *HvSCAR*-*A*, *HvSCAR*-*B*, or *HvSCAR*-*C* in *M. truncatula* A17 *API* and *api* hairy roots. The same vector backbone carried a DsRed expression cassette, which serves as a fluorescent reporter of successful root transformation. Expression of *HvSCAR*-*B* and *HvSCAR*-*C* restored *api* root hair length to A17 control levels, whereas *HvSCAR*-*A* expressing *api* root hairs were significantly shorter than A17 and statistically indistinguishable from *api* transformed with an empty-vector control (Fig. 1b,c). Thus, the barley genes *HvSCAR-B* and *HvSCAR-C* (CII-clade) can functionally replace *MtAPI* in *M. truncatula* roots, whereas *HvSCAR-A* (CI-clade) cannot. This suggests that while functional divergence occurred after the initial CI/CII clade split, the duplicated genes *HvSCAR-B* and *HvSCAR-C* both retained the ability to regulate actin dynamics during root hair growth.

### CRISPR-CAS9 knockout of individual barley *HvSCAR* genes does not affect vegetative growth but impairs seed production in *hvscar-a*

*M. truncatula api* mutants show impaired root-hair development but otherwise produce healthy plants without macroscopic defects in growth or seed production (Gavrin et al., 2020). To assess whether loss of *HvSCAR* function affects development, we generated CRISPR-Cas9 knockouts using two guide RNAs per gene (Figure S4). From the initial transformations, we recovered transformants for all three genes and advanced at least two confirmed mutant lines per gene to the T₃ generation to obtain *Cas9*-free, homozygous knockouts.

Individual T_1_ generation *hvscar*-*a* plants from independent parents produced significantly fewer seeds than *hvscar*-*b, hvscar*-*c*, and Golden Promise (GP) controls but otherwise seemed to behave normal (Figure S5a,b,d,e). To validate this observation, we simultaneously grew six homozygous, T5 generation plants from one confirmed knockout line per gene under identical conditions. Vegetative growth was comparable among Golden Promise and *hvscar*-*a*, *hvscar*-*b*, and *hvscar*-*c* single mutants up to seed development (Figure 2a). Consistent with the T1-generation results, *hvscar*-*a* plants produced significantly fewer total seeds per plant (Figure 2d) due to a significant reduction in seed production per spike (Figure 2e) while *hvscar-b* and *hvscar-c* plants were not impaired. Unlike the shortened *M. truncatula api* root hairs, none of the single *hvscar* knockout mutants showed reduced root hair length (Figure 2f).

Taken together, these results indicate that individual *HvSCAR* genes execute distinct, non-redundant functions in barley development. While none of the single mutants exhibited defects in vegetative growth or root hair elongation, *HvSCAR*-*A* is specifically required for normal seed set, as its loss reduces seed formation per spike. By contrast, *hvscar*-*b* and *hvscar*-*c* do not differ from Golden Promise in these seed-related traits.

**Figure 2:**
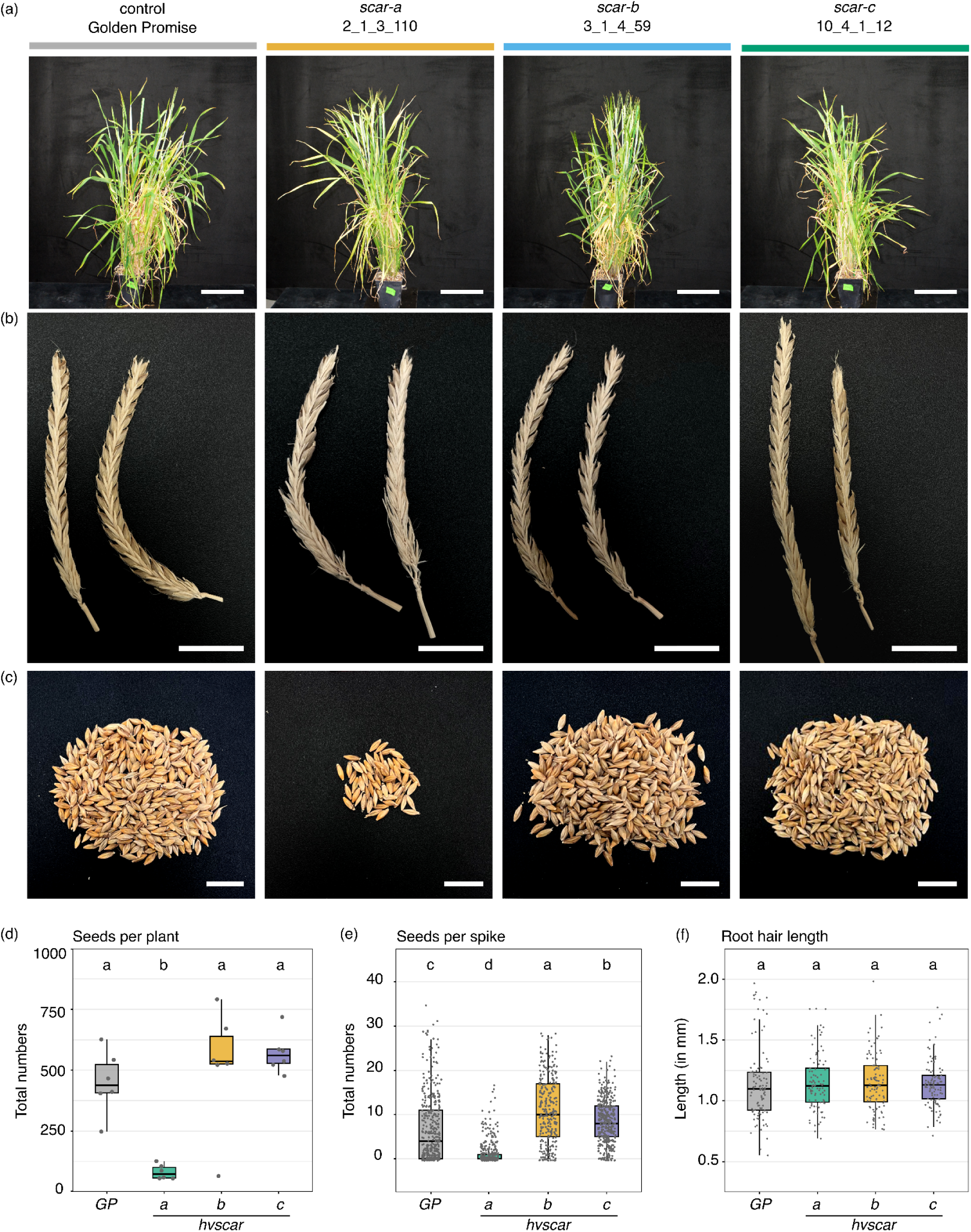
Development and seed set analysis of single *hvscar* CRISPR knockout mutants. (a) Representative images of 9-week-old Golden Promise and *hvscar* single CRISPR knockout plants grown under identical conditions. Scale bar: 20 cm. (b) Images of two representative spikes for each genotype. Awns were manually removed from spikes to improve visualization of the seeds. Scale bar: 5 cm. (c) Images of a representative batch of seeds harvested from one individual plant per genotype. Scale bar: 2 cm. (d) Number of seeds per plant: Each data point represents the total number of seeds for one individual plant (n =6 for each genotype). (e) Number of seeds per spike: Each data point represents the number of seeds that were harvested from one spike. The data for all spikes from 6 different plants for each genotype are shown (*GP*, n = 403; *hvscar-a*, n = 372; *hvscar-b*, n = 285; *hvscar-c*, n= 404). (f) Root hair length measurements: The length of 20 root hairs, measured in mm, was analysed for 5 roots from different plants for each genotype. Each datapoint represents one measurement (n = 100 for each genotype). Statistics: The data was analysed for normality with a Shapiro-Wilk test, followed by a Kruskal-Wallis with Bonferroni correction (d, e, f); statistical significance groupings are indicated by letters.

### *hvscar-b,c* plants have shorter root hairs but vegetative growth and seed development are not impacted

Since both *HvSCAR-B* and *HvSCAR-C* complemented the *M. truncatula api* root hair phenotype, we hypothesized that these genes may function redundantly during barley root hair development. To test this, we generated an *hvscar-b,c* double mutant by crossing *hvscar-b* line 3-1 with *hvscar-c* line 10-4 and selected an azygous *HvSCAR-B,C* control line from the same cross (Figure S4b,c). Root hair length measurements in F5 seedlings revealed that *hvscar-b,c* developed significantly shorter root hairs than the *HvSCAR-B,C* control (Figure 3a-b). In contrast, overall plant morphology was indistinguishable between *hvscar-b,c* and control plants (Figure 3c-e). Neither the total seed number per plant nor the number of seeds per spike differed significantly from controls in the F5 and the previous F4 generation (Figure 3f,g and Figure S5c,f). These plants were grown alongside the corresponding single mutants and Golden Promise plants shown in Figure 2, and no significant differences were detected between Golden Promise and the azygous *HvSCAR-B,C* line.

Together, these findings indicate partial functional redundancy between *HvSCAR-B* and *HvSCAR-C* in barley root hair elongation. While simultaneous loss of both genes reduces root hair length, vegetative development and overall reproductive performance remain largely unaffected under the tested growth conditions.

**Figure 3:**
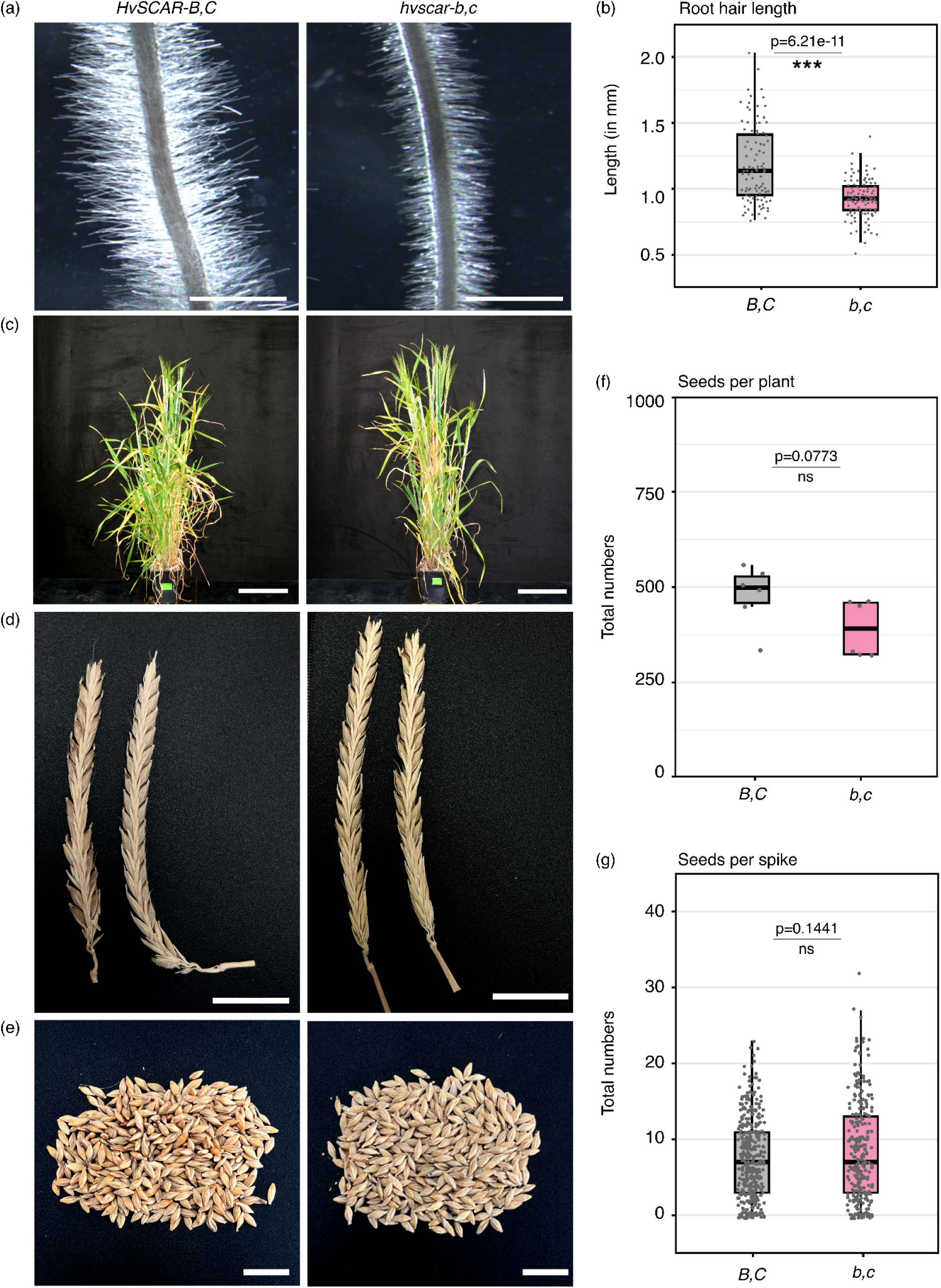
Development and seed set analysis of *hvscar*-*b,c* and corresponding azygous plants. (a) Representative images of developmentally comparable root hair zones of *HvSCAR-B,C* and *hvscar-b,c* roots. Scale bar: 2 mm. (b) Root hair length measurements. The length of 20 root hairs, measured in mm, was analysed for 5 different roots for each genotype. Each datapoint represents one measurement (n = 100). (c) Representative images of 9-week-old *HvSCAR-B,C* and *hvscar-b,c* plants grown under identical conditions (grown at the same times with plants from Figure 2). Scale bar: 20 cm. (d) Images of two representative spikes for each genotype. Awns were manually removed from spikes to improve visualization of the seeds. Scale bar: 5 cm. (e) Images of a representative batch of seeds harvested from one individual plant per genotype. Scale bar: 2 cm. (f) Number of seeds per plant: Each data point represents the total number of seeds for one individual plant (n = 6 for each genotype). (g) Number of seeds per spike: Each data point represents the number of seeds that were harvested from one spike. The data for all spikes from 6 different plants for each genotype are shown (*HvSCAR-B,C*, n = 391; *hvscar-b,c*, n = 281). Data normality was assessed using the Shapiro–Wilk test. Depending on the outcome, either a Mann-Whitney-U test (b, g) or a Welch’s *t*-test (f) was applied. Corresponding *p*-values are shown in the panels. Significance levels are indicated as ns (p ≥ 0.05) and *** (p < 0.001).

### Interactions of the *hvscar-b,c* mutant roots with detrimental and beneficial filamentous microbes are differentially affected

In *M. truncatula*, the *api* mutant shows increased resistance to *P. palmivora* infections (Gavrin et al., 2020). Because the barley *hvscar-b,c* mutant shares the same reduced root hair phenotype as *api*, we investigated whether these homologs also provide enhanced resistance against the same pathogen.

We employed the *P. palmivora* strain FLIMA for infection assays because previous tests identified FLIMA as one of the most aggressive isolates on barley cv. Baronesse roots (Le Fevre et al., 2016). We transformed FLIMA with the red fluorescent marker tdTomato (FLIMA-td) to enable infection visualization and detailed observation of colonization structures. At three days post-inoculation (dpi), we detected patchy fluorescence along the roots of both *HvSCAR*-*B,C* and *hvscar*-*b,c* plants (Figure S6a), and by six dpi strong sporulation was evident near the root tips (Figure S6b). However, visually we did not observe clear qualitative differences in infection levels between the two genotypes. To assess potential quantitative differences in pathogen biomass, we performed qRT-PCR at 3 dpi using the *P. palmivora* biomass marker *PpEF1α*. On average, *hvscar*-*b,c* roots displayed ∼50% lower *PpEF1α* expression than wild-type roots, indicating reduced pathogen biomass (Figure 4a). We also observed a significant reduction in expression of the sporulation marker *PpCdc14* relative to barley reference genes, but not FLIMA reference genes, consistent with lower biomass resulting in reduced sporulation (Figure 4b and c).

Comparing root expression patterns using RNAseq between control and *hvscar-b/c* double mutant returned a high level of correlation (R² value of 0.962) with only 80 differentially expressed genes (log₂FC ≥ |2| and *padj* < 0.05) between the genotypes (Figure S7 and Data S3). This suggests that roots of *hvscar-b/c* double mutants do not mount a prominent constitutive defense response in absence of an infection.

We then examined whether the *hvscar*-*b,c* mutations also affect interactions with beneficial filamentous microbes. Arbuscular mycorrhizal (AM) fungi, like oomycetes, establish intimate associations within root tissues that involve host cell remodeling and the formation of intracellular colonization structures. Upon inoculation with *F. mosseae*, *hvscar*-*b,c* roots exhibited significantly increased colonization compared to wild-type roots, with an average enhancement of ∼15% (Figure 4g). In contrast, key symbiotic structures like arbuscules and vesicles were present at comparable frequencies in both genotypes (Figure 4h,i), indicating that the *hvscar*-*b,c* mutations facilitate the entry and/or spread of the fungus within the root system without altering its intracellular developmental program.

Together, these results demonstrate that *hvscar*-*b,c* mutations enhance resistance to the oomycete pathogen *P. palmivora* without inducing constitutive immune activation. Remarkably, these same mutations promote, rather than impair, colonization by the beneficial AM fungus *F. mosseae*. Thus, loss of specific *HvSCAR* genes selectively rewires root - microbe interactions.

**Figure 4:**
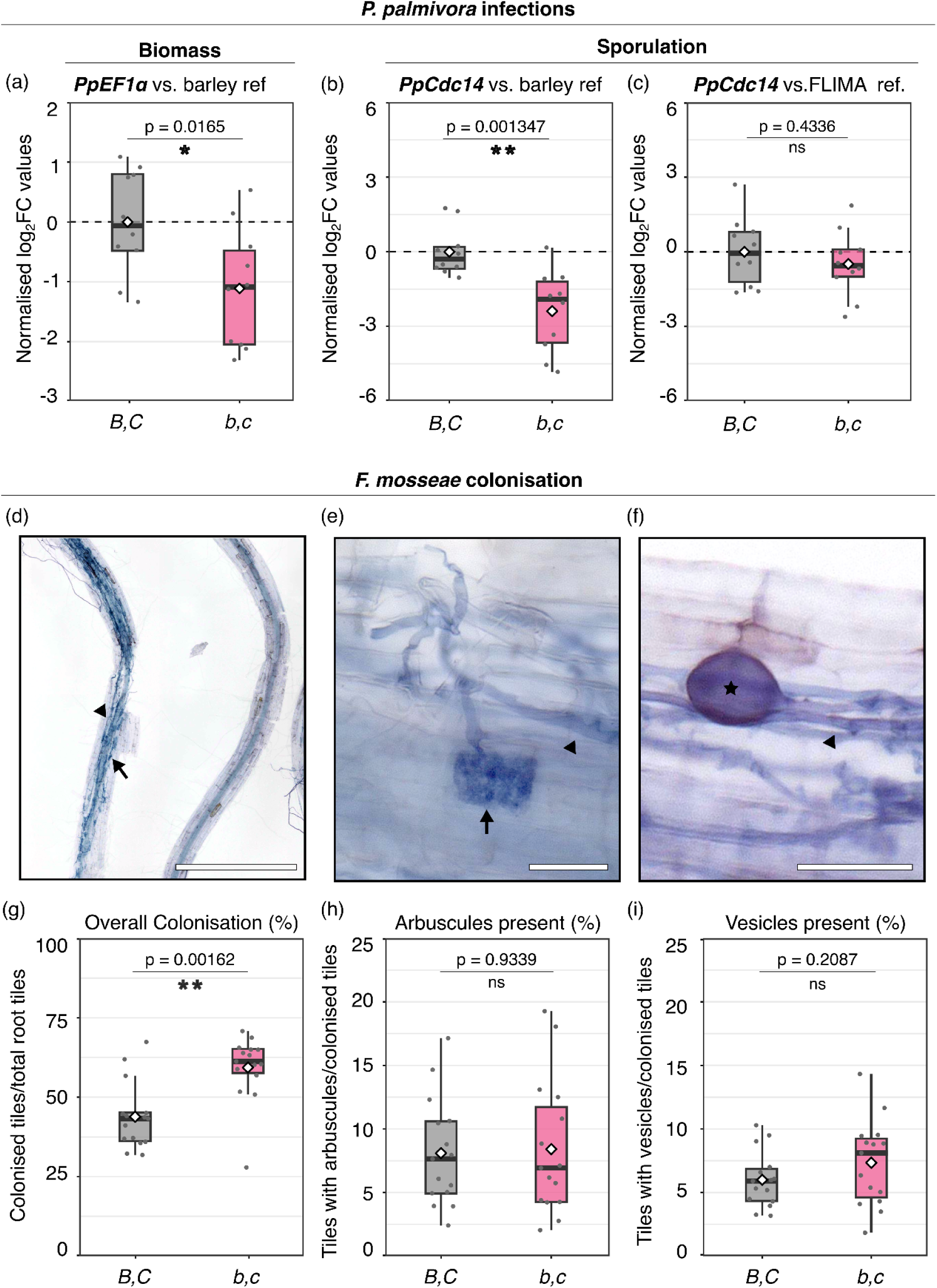
Root pathogenic *Phytophthora* and symbiotic fungi are oppositely impacted in their colonisation of hv*scar-b,c* plants. (a-c) Normalised log₂ fold-change (log₂FC) in expression of *P. palmivora* biomass markers genes *PpEF1α* (a) and *PpCdc14* (b), calculated from efficiency-corrected ΔΔCq ratios with either *HvEF1α* and *HvCyc* as barley reference genes (a,b) or *PpEF1α* and *PpWS21* as FLIMA reference genes (c). Expression values were log₂-transformed and normalised to the mean expression of the wild-type genotype (*B,C*), set to 0 (dashed line). Boxplots display the distribution of normalised log₂FC values across genotypes with white diamonds indicating the means. Individual points represent the mean of three technical replicates per biological sample (n = 8). (d-f) Representative images of barley root fragments partially colonised by arbuscular mycorrhiza (d), with close-ups of fungal structures including hyphae (black triangles), arbuscules (e, black arrow), and vesicles (f, black star). (g-i) Boxplots show overall colonisation levels (g), quantification of arbuscules (h) and vesicles (i). Individual points represent ratios from analysed microscopy slides of root fragments from individual plants (n = 15). Statistics: Data normality was assessed using the Shapiro–Wilk test. Depending on the results, either Welch’s *t*-test (a,b,c,i) or the Mann-Whitney-U test (g, h) was applied. Corresponding *p*-values are shown in the panels. Significance levels are indicated as ns (p ≥ 0.05), * (p < 0.05), and ** (p < 0.01).

## Discussion

Soil-borne pathogens threaten global crop production, and their impact is expected to intensify as climate change alters pathogen ranges and soil microbiome dynamics. Because roots remain difficult to monitor and protect, strategies that disable host susceptibility (S) factors represent an attractive complement to classical, race-specific *R*-gene approaches. Yet actionable *S*-genes for cereal root diseases are largely unknown. Here, we show that a role for *SCAR* genes in root susceptibility originally discovered in dicots, is conserved in the monocot barley. Under laboratory conditions, loss of the barley *SCAR* homologs *HvSCAR*-*B* and *HvSCAR*-*C* enhances quantitative resistance to *P. palmivora* without major penalties on plant development or AMF symbiosis.

### Identification and functional diversification of barley SCARs

We performed a targeted phylogenetic analysis to identify bona fide barley SCAR/WAVE proteins relevant to MtAPI function. We focused on proteins with canonical SCAR domain architecture and manually curated all candidates, an approach that likely excluded monocot WASH proteins whose structure differs markedly from SCAR/WAVE proteins (Veltman and Insall, 2010) and therefore lay outside our analytical scope. As our goal was to resolve barley HvSCAR proteins rather than all ARP2/3 activators, we restricted the phylogeny accordingly.

Our data reveal clear functional divergence between barley *HvSCARs* of clade I and clade II. The clade I gene *HvSCAR*-*A* is required for normal reproductive development, as *hvscar*-*a* mutants show a strong reduction in seed set. A reduced reproductive output has also been reported for the rice clade I *SCAR* mutant *tut1*, although in rice this results from defects in panicle development rather than seed formation (Bai et al., 2015). Meanwhile, *hvscar*-*b*, *hvscar*-*c*, and the *hvscar*-*b,c* double mutant do not affect seed production. Comparisons with other species indicate that *SCAR* paralog functions vary across lineages: in *A. thaliana*, several clade I and II *AtSCARs* show substantial redundancy (Zhang et al., 2008), particularly in trichome development, whereas in *M. truncatula*, the clade II genes *MtAPI* and *MtHAPI1* have separable functions in root hair formation (Brumm et al., 2025). These cross-species differences indicate that *SCAR* paralogs can adopt redundant or specialized roles depending on the plant lineage. In barley, this separation offers practical advantages: distinct *SCAR* functions enable targeted modification of clade II genes to improve root-disease resistance without compromising reproductive performance.

### Context**-**dependent microbial responses in *HvSCAR***-***B,C* mutants with limited developmental costs

Because susceptibility-gene modifications can cause unintended growth or yield penalties (van Schie and Takken, 2014), it is notable that the *hvscar*-*b,c* double mutant showed no major vegetative defects under our laboratory conditions. The plants had shorter root hairs and reduced spike formation but normal overall seed production. The widely employed *mlo* mutation, which provides strong resistance against powdery mildew infections in cereals, can lead to lesion mimicry and correlated yield costs in the field (Kjær et al., 1990). Whether *hvscar*-*b,c* mutations, or their associated root hair phenotype, affect reproductive performance under field conditions remains to be tested. Mechanistically, the enhanced quantitative resistance of *hvscar*-*b,c* to *P. palmivora* is unlikely to arise from constitutive immune activation, as RNA-seq showed no broad immune priming. More plausibly, loss of *HvSCAR*-*B* and *HvSCAR*-*C* may alter cell wall deposition or plant actin-dependent microbe entry processes. While *M. truncatula api* mutants exhibit modified pectin and xyloglucan compositions (Gavrin et al., 2020), barley’s Type II cell walls rely on different polymers (e.g., glucuronoarabinoxylans and mixed linkage glucans) (Carpita and Gibeaut, 1993; Kim and Brandizzi, 2021). If resistance in *M. truncatula* and barley is secretion-driven, the shared trait likely involves the actin-dependent secretory hub itself rather than a single conserved cell wall polymer. Future cell wall fractionation and transport assays will be needed to distinguish these possibilities. Interestingly, *hvscar*-*b,c* roots also supported a greater extent of AM colonisation, while maintaining normal arbuscule and vesicle formation, indicating that SCAR-dependent trafficking influences both pathogenic and symbiotic entry. Each type of microbial interaction was assessed in its relevant biological context: *P. palmivora* infection on young elongating seedling roots, and AM symbiosis in mature differentiated roots. These developmental stages naturally differ in wall composition and secretory activity, which may contribute to the contrasting outcomes. Age-matched infection assays and zone-resolved imaging could help clarify how *SCAR*-dependent processes shape susceptibility across specific tissues and developmental stages.

### Implications for disease resistance engineering in cereals

Our findings provide a proof-of-concept for root-targeted S-gene engineering in cereals. The *hvscar*-*b,c* mutant combines quantitative seedling resistance to *P. palmivora* with normal growth and enhanced AMF colonisation, highlighting *SCAR* gene modification as a promising plant engineering strategy. Future studies should evaluate the susceptibility spectrum of *hvscar-b* and *hvscar-c* mutants against a diverse panel of root- and shoot-infecting oomycete and fungal pathogens representing different lifestyles. Field trials will be essential to evaluate whether *hvscar*-*b,c* confers stable resistance and maintains yield across soils, climates, and microbiomes. Removal of *MtAPI* in *M. truncatula* for example leads to a significant increase in the root associated microbiota diversity (Tkacz et al., 2022). If agronomically relevant penalties occur under field trials, targeted approaches such as tissue-specific genome editing or tuned expression via promoter lesions could strike a balance between quantitative resistance and developmental needs. Taken together, our results position clade II *SCAR/WAVE* genes as actionable susceptibility factors for enhancing root health in cereals. As soil-borne disease pressure intensifies, targeted modulation of *HvSCAR-B* and *HvSCAR-C* may provide a compatible and complementary route to broaden pathogen protection alongside established R-gene based strategies.

### Experimental procedures

#### Phylogenetic Analysis

SCAR protein sequences were obtained from public genome and proteome databases (Data S1). Sequences were aligned to *M. truncatula* and *A. thaliana* SCARs using MUSCLE (CLC Main Workbench 20.0.4) to assess SHD/VCA domain presence/absence. Full-length sequences were aligned and subsequently automatically trimmed using the bioconda packages “mafft” (version 7.471; (Rozewicki et al., 2019)) and “trimAI” (version 1.4.1; (Capella-Gutierrez et al., 2009)). Phylogenetic trees were constructed using the bioconda package “IQ-TREE” (version 2.0.3; (Nguyen et al., 2015)) with maximum likelihood and bootstrapping (1000Ufboot+SH-aLRT) and identified the best fitting substitution models: “model VT+F+I+G4” for Figure 1a and “JTT+I+G4” for Figure S1. Trees were visualised with the online tool “interactive Tree Of Life” (iTOL version 6.9.1; (Letunic and Bork, 2024)). The trees were rooted at the midpoint and branches with less than 50% SH-aLRT support (bootstrap 0) were deleted for Figure 1 and Figure S1. The trees were stylistically modified for figure presentation using Inkscape (version 1.2.2).

#### Construct design and cloning

Primer design, sequence assembly, and analyses were conducted using CLC Main Workbench 20 (Qiagen). Primer sequences are listed in Table S1. The pKGW-MGW-pAPI:API construct was kindly provided by Aleksandr Gavrin (Gavrin et al., 2020). The *HvSCAR-B* (*HORVU.GOLDEN_PROMISE.PROJ.4HG00344290*) coding sequence (CDS) was synthesized by GenScript in pUC57 with flanking attL1/attL2 sites, based on the Golden Promise Pangenome v2 annotation, identical to Morex v2 (2019) and v3 (2021) annotation. The *HvSCAR-A* (*HORVU.GOLDEN_PROMISE.PROJ.3HG00222400*) CDS was amplified from

Golden Promise root cDNA using primers SB103/SB104 and Phusion™ High-Fidelity DNA Polymerase (NEB) with GC buffer. Sequence analysis revealed an additional CAG at positions 720-722, adding one serine at amino acid 240; this triplet is present in Morex v1 and v3, but absent from Morex v2 and Golden Promise v2. Subsequent constructs used the verified Golden Promise sequence. *HvSCAR-C* (*HORVU.GOLDEN_PROMISE.PROJ.5HG00469460*) CDS was amplified from 9-day-old seedling shoot cDNA using primers SB101/SB102 and Phusion™ polymerase with GC buffer. Comparison with the Golden Promise v2 annotation revealed an 87 bp insertion (CDS positions 6620–6706), absent from Morex v2 and v3 annotations. Morex v1 is incomplete (567bp). All constructs used the verified Golden Promise sequence.

*HvSCAR-A* and *HvSCAR-C* CDS PCR products were cloned into pDONR221 via Gateway® BP Clonase™. Initial pDONR-SCARC clones contained nonsynonymous base-pair substitutions; to obtain a mutation-free construct, a 2082-bp fragment from pDONR-SCARC #3 (NdeI/ScaI-HF) was ligated into the 7131-bp backbone of pDONR-SCARC #23 (NdeI/ScaI-HF digested, Antarctic Phosphatase-treated), generating pDONR-SCARC #3-23-10. Both #3 and #3-23-10 were digested with EcoRI/HindIII, and a 2934-bp fragment from #3 was ligated into the 6285-bp backbone of #3-23-10, yielding the final mutation-free clone pDONR-SCARC #3-23-10-3-1 (SB95_pDONR_HvSCARC).

All entry and destination vectors were verified by diagnostic restriction digest and Sanger sequencing (Source BioScience). Destination clones were assembled by recombining SB88_pDONR_HvSCARA, SB98_pUC57(Amp)_HvSCAR-B, and SB95_pDONR_HvSCARC with pENTR4_1_prMtAPI and pENTR_p2rp3_T35STerm into pKGW-RR-MGW using LR Clonase® Plus (Thermo Fisher Scientific). Primer sequences and constructs are in Tables S1 and S2; vector and CDS sequences are in Data S2.

Level 1 sgRNA expression constructs were generated by amplifying the sgRNA scaffold from pICSL90010 using: for sgRNA1, HvSCAR-A (SB93/SB90), HvSCAR-B (SB88/SB90), HvSCAR-C (SB99/SB90); for sgRNA2, HvSCAR-A (SB107/SB90), HvSCAR-B (SB108/SB90), HvSCAR-C (SB100/SB90). sgRNA1 amplicons were assembled with the Level 0 Triticum aestivum U6 promoter (pICSL90003) and Level 1 position-3 acceptor vector pICH47751 via MoClo. sgRNA2 amplicons were assembled with pICSL90003 and Level 1 position-4 acceptor vector pICH47761. Binary plasmids were assembled using Golden Gate MoClo (Weber et al., 2011). Final constructs combined the Level-M backbone pAGM8031, Level-M end-linker pICH50900, position 1 35S::hptII::35S hygromycin cassette (pICSL11059), position 2 Zm::Cas9::nos cassette (pICSL11056), position 3 sgRNA1 module, and position 4 sgRNA2 module (Data S2).

#### Plant materials and growth conditions

The two-rowed *Hordeum vulgare* accession Golden Promise (PI 343079) was obtained from the USDA Germplasm Resources Information Network (Aberdeen, ID, USA) (Bettgenhaeuser et al., 2021). Following tissue-culture regeneration, T₀ plants of the CRISPR lines were transferred to John Innes Cereal Mix and grown in greenhouses under 18°C day (16 h) and 12°C night (8 h) conditions with supplemental illumination from 400 W HQI metal halide lamps. For subsequent generations, barley seeds were stratified for two to three days in the dark at 4°C and then germinated for two to three days on wet Whatman filter paper in the light at 24°C. Seedlings were transplanted into a 1:1 mixture of Levington F2 compost and vermiculite and maintained in a Conviron growth chamber with a 16-h light/8-h dark cycle, day/night temperatures of 18°C and 15°C, 80% relative humidity, and a light intensity of approximately 500 μmol m⁻² s⁻¹. For mycorrhiza assays, seedlings were grown in a sand/terragreen mix substrate in a Conviron growth chamber set to a 16-h light/8-h dark cycle at 21°C.

#### Generation of transgenic *Medicago truncatula* roots and root hair analysis

The destination constructs were transferred into *Agrobacterium rhizogenes* strain *Arqua1193* (rifampicin and carbenicillin resistant) and transformed into the roots of *M. truncatula* A17 and *api* seedlings (Limpens et al., 2005). Following transformation, roots were maintained on Fahraeus medium for a duration of three weeks. Three weeks after transformation, epifluorescence images of *M. truncatula* roots expressing DsRed were acquired using a Leica M165 FC Fluorescent Stereomicroscope equipped with a DFC310FX camera and DSR filter (10447412). All images were captured at a consistent magnification, including a reference scale, using Leica Application Suite Software (version 4.8.0). Root hair length was measured in ImageJ2 using the ROI Manager and Freehand Line tools. Image scales were globally calibrated via the SetScale function. For each genotype, 10 root hairs were measured from five independently transformed roots (n = 50), and lengths were recorded in millimetres. To account for variation in imaging depth, root hair length (in mm) was normalized to the diameter of the respective root by calculating root hair length/diameter ratios. Data distribution was evaluated using the Shapiro-Wilk test. When p-values were below 0.5, non-parametric comparisons were performed using the Kruskal-Wallis test followed by Bonferroni-adjusted post hoc analysis (α = 0.05). Box plots were generated in R and adapted to fit the figure panel in Incscape. Images were processed for brightness, and contrast adjustments using GIMP. Final figure panels were assembled in Inkscape, with scale bars manually added based on the reference scale. Each construct was analysed in two independent transformation experiments, with one representative dataset selected for figure presentation.

#### Generation of Hordeum vulgare CRISPR knockout mutants

For each HvSCAR gene, two 20-nt target sequences immediately upstream of a PAM were selected using CRISPOR (https://crispor.gi.ucsc.edu/). One gRNA was designed in the first exon and the second in the largest exon (*HvSCAR*-*A*: exon 6; *HvSCAR*-*B*: exon 6; *HvSCAR*-*C*: exon 8). Guides were retained only if they had a single predicted 20-mer target site in the barley genome and included a nearby restriction site: *HvSCAR*-*A* guide 1: ScaI; guide 2: AccI; *HvSCAR*-*B* guide 1: SmaI/XmaI; guide 2: BsrGI; *HvSCAR*-*C* guide 1: HaeII; guide 2: NspI. CRISPR/SpCas9 vectors carrying the *HvSCAR*-targeting gRNAs were electroporated into *Agrobacterium tumefaciens* strain AGL1 (Gene Pulser, Bio-Rad) following (Lazo et al., 1991), and used to generate transgenic spring barley cv. ‘Golden Promise’ (Hensel et al., 2009). Single knockout lines were identified by PCR and Sanger sequencing of the targeted genomic regions. Genotyping primers were: *HvSCAR*-*A* guide 1: SB207/SB162; guide 2: SB160/SB184; *HvSCAR*-*B* guide 1: SB180/SB166; guide 2: SB85/SB181; full HvSCAR-B deletion: SB180/SB181; *HvSCAR*-*C* guide 1: SB185/SB186; guide 2: SB209/SB187. Cas9 presence was assessed with SB233/SB234 (see Table S1).

From an initial screen of 42 *HvSCAR*-*A*, 78 *HvSCAR*-*B*, and 50 *HvSCAR*-*C* T₀ transformants, at least three mutation-positive T₁ lines per gene were selected (*hvscar-a*: SB47-2-1, SB47-10-3, SB47-5-4, SB47-5-5; *hvscar-b*: SB46-3-1, SB46-24-1, SB46-27-1; *hvscar-c*: SB48-10-4, SB48-12-2, SB48-16-2). These were advanced to T₂ and analyzed for seed formation (SB47-5-5: only 186 seeds from four segregants). Two representative lines per gene were propagated to T₅ to obtain Cas9-free, homozygous plants (*hvscar-a*: SB47-2-1-3-110, SB47-5-5-5-220; *hvscar-b*: SB46-3-1-2-59, SB46-24-1-2-33; *hvscar-c*: SB48-10-4-1-12, SB48-16-2-1-84) and seed numbers were counted for one representative line per gene.

To generate the *hvscar*-*b,c* double mutant, the Cas9-free, homozygous *hvscar-b* line SB46_3_1_2_59 and *hvscar-c* line SB48_10_4_1_12 were crossed. Double-homozygous F₂ segregants were identified by genotyping and confirmed by Sanger sequencing. Seed numbers were quantified in the F₄ and F₅ generations.

#### Root hair and developmental phenotyping of *H. vulgare* CRISPR mutants

Seeds were germinated for 48 h on sterile Whatman paper at 24°C, transferred to ½ basal medium (5% sucrose) and grown for 4 days at 20°C before imaging with a Leica DMi8 fluorescence microscope. Root hair length was quantified in ImageJ2 using the ROI Manager. Roots were straightened, and hairs were measured with the freehand line tool within a defined region corresponding to 25–30% of the total root length from the tip. This ROI was divided into 10 equal sub-zones, and one hair per sub-zone per side was measured. Image scales were globally calibrated using SetScale. Five roots per genotype were analysed, with 20 hairs measured per root. For vegetative growth assessment, representative T5 and F5 plants were photographed 9 weeks after germination using a Nikon D5200 with a Nikkor 18–55 mm f/3.5–5.6G VR lens; a meter scale was included for size reference. At maturity, all spikes per plant were harvested, and the number of seeds per spike were manually counted. Data were analysed and plotted in R, and plots were formatted in Inkscape. Total seed number per plant was calculated as the sum of all seeds across spikes. Representative spikes and seeds were photographed after harvest using an iPhone 13 with a ruler for scale; awns were removed to improve visibility. Images were cropped and adjusted for brightness and contrast in GIMP, and final figure panels were assembled in Inkscape with manually added scale bars.

#### *P. palmivora* transformation and barley root infection assays

Generation of the fluorescent *P. palmivora*strain FLIMA-td involved electroporation-based transformation of FLIMA (accession number P7545, derived from the Philippines) zoospores with a pTOR::tdTomato expression construct, as described for the ARI-td strain previously (Le Fevre et al., 2016). For qRT-PCR analysis, barley root infection assays with FLIMA-td were performed as described before (Macleod et al., 2025). Briefly, barley seeds were surface-sterilized in 70% ethanol (5 min) and 2% sodium hypochlorite (5 min), rinsed once between steps and at least four times afterward, then placed on wet Whatman paper at 4°C in the dark for 2–3 days and germinated at 24°C in the dark for 4 days. Seedlings were transferred to 1% phytoagar in 100-mm square plates (up to 12 per plate), and roots were covered with 80-mm 325P cellulose discs. Each root system received 100 μL of FLIMA-td zoospores (5 × 10⁴ mL⁻¹). Plates were wrapped at the base in black foil, incubated horizontally at 25°C under constant light to allow spore attachment, then tilted 80° to prevent root growth into the medium. For qRT-PCR, roots were harvested 3 dpi, washed, and flash-frozen.

For imaging, seedlings were placed on 1% phytoagar 3 days after germination. Plates were cast with a thinner lower half to form a basin for roots, which were submerged in 20 mL of FLIMA-td zoospores (5 × 10⁴ mL⁻¹). Plates were wrapped at the base in black foil and incubated horizontally at 25°C. Whole-root imaging at 3 and 6 dpi was performed using a Leica M205 FCA stereomicroscope with a Leica K5-14403327 camera (1.0×/0.03 NA, 1.39× zoom). Tile scans were acquired after removing the plate lid. Images were processed using Thunder Image Clearing, with mCherry used to detect FLIMA-td fluorescence and a GFP filter to visualize root autofluorescence. Channels were merged in ImageJ.

#### qRT-PCR

For qRT-PCR analysis of *FLIMA*-*td* infected roots, one barley root system was used per sample. Total RNA was extracted using the Spectrum Plant Total RNA Kit (Sigma-Aldrich) followed by on-column DNase treatment with the RNase-Free DNase Set (Qiagen). cDNA was synthesized from 1 µg of total RNA using the Roche Transcriptor First Strand cDNA Synthesis Kit. qPCR reactions were prepared with the Roche LightCycler 480 SYBR Green I Master Mix and run in 10-µl triplicate reactions on a Roche LightCycler 480 II instrument. The cycling program consisted of an initial denaturation at 95 °C for 5 min, followed by 45 cycles of 95 °C for 10 s, 60 °C for 10 s, and 72 °C for 10 s. Primer sequences are listed in Table S1. Gene expression was quantified in R using the 2^−ΔCt method, calculating *PpEF1α* relative to *HvEF1α* and *HvCyc* (as a relative measure of pathogen biomass), or *PpCdc14* relative to *PpEF1α* and *PpWS21* (as an indicator of sporulation) (Le Fevre et al., 2016). The results were subsequently plotted in R.

#### *F. mosseae* root colonisation assays

Three-day-old seedlings were transferred to pots containing 500 mL of a sand:terragreen mixture supplemented at a 1:10 ratio with *Funneliformis mosseae* crude inoculum (MycAgro, France) and watered with deionized (DI) water. Plants were kept in a plastic tent for one week to maintain high humidity, then grown at 24 °C under a 16 h light/8 h dark photoperiod. After two weeks, plants were supplied with Hoagland nutrient solution (per litre: 4 mL 5 mM KH₂PO₄ and 0.75 g Hoagland’s No. 2 Basal Salt Mixture, Modification 2 without ammonium phosphate; Caisson Labs). After eight weeks, whole root systems were washed free of substrate and three root segments (upper, middle, lower) were collected per plant. Fungal structures were visualized using a modified ink–vinegar staining protocol (Vierheilig et al., 1998). Roots were cleared in 10% (w/v) KOH for 10 min at 95 °C, rinsed in 5% (v/v) acetic acid, and stained in 5% Sheaffer black ink/5% acetic acid for 10 min at 95 °C. Stained root fragments were rinsed in DI water, briefly cleared in ClearSee (30 seconds per sample; (Kurihara et al., 2015)), and placed on a microscopy slide in mounting media (20% glycerol, 50 mM Tris-HCl pH 7.5, 0.1% Tween-20). Tile-scanned images were acquired at 200× magnification using the Keyence microscope VHX 7000 (2D stitching mode) and assembled using the Fiji Grid/Collection Stitching plugin. White levels and artefacts (air bubbles) were manually adjusted/removed from the images in GIMP to simplify automated analysis. Total colonisation was quantified using AMFinder software (Evangelisti et al., 2021) with the CCN1 classifier, and arbuscule and vesicle structures were quantified using the CCN2 classifier.

#### RNA sequencing analysis

The roots of 3 *HvSCAR-B,C* and 5 *hvscar-b,c* seedlings, grown on 1% Phytoagar, were harvested 8-days after germination and RNA was extracted as described for qRT-PCR analysis. RNA samples were sequenced by Novogene UK using a paired-end strategy (2 × 150 bp), yielding approximately 40 million reads per sample. Read quality was assessed using FastQC. Reads were mapped to the annotated BPGv2 Golden Promise barley assembly (release 220214), publicly available via the IPK Gatersleben Galaxy server (https://galaxy-web.ipk-gatersleben.de/libraries/folders/Fd071e794759ab192; (Jayakodi et al., 2024)). The nf-core/rnaseq workflow (version 3.9; (Ewels et al., 2020)) was used for read alignment and quantification, implementing STAR for mapping (Dobin et al., 2013) and Salmon for transcript-level quantification (Patro et al., 2017).

Gene-level count tables were imported into R, rounded to integer values, and prefiltered to remove lowly expressed genes by retaining only genes with a summed read count greater than one across all samples. Filtered counts were normalized using DESeq2 (version 1.42.2; (Love et al., 2014)) through estimation of size factors to correct for differences in sequencing depth. Differential expression analysis was performed using DESeq2, comparing *hvscar-b,c* samples against *HvSCAR-B,C* control samples. Genes with an adjusted p-value (padj) < 0.05 and an absolute log2 fold change (|log2FC|) ≥ 2 were considered differentially expressed. A complete list of identified differentially expressed genes (DEGs) and their transcript per million (TPM) values is provided in Data S2.

### Statistical analysis

All statistical analyses were conducted in R (version 4.3.3). Data were tested for normality using the Shapiro–Wilk test. Depending on the outcome, appropriate single- or multiple-comparison post hoc tests were applied. The specific statistical test used for each analysis is indicated in the corresponding figure legend, and exact p-values or letters indicative of significance groupings are reported within the figure panels.

## Accession numbers

*Hv*SCAR-A (HORVU.GOLDEN_PROMISE.PROJ.3HG00222400), *Hv*SCAR-B (HORVU.GOLDEN_PROMISE.PROJ.4HG00344290), *HvSCAR-C* (HORVU.GOLDEN_PROMISE.PROJ.5HG00469460), *Mt*API (Medtr4g013235/MtrunA17_Chr4g0004861)

## Author Contributions

Conceptualization: S.B. and S.S. Methodology: S.B., Ma.Ma., I.H.P., E.E., Ma.Mü., M.J.M., S.S. Investigation: S.B., Ma.Ma., I.C., I.H.P. Validation: S.B., Ma.Ma., I.C., I.H.P. Formal analysis: S.B., Ma.Ma. Data curation: S.B. Visualization: S.B. Supervision: S.B. and S.S. Project administration: S.S. and S.B. Writing - original draft: S.B. Writing - review and editing: S.B. Ma.Ma., E.E., Ma.Mü., M.J.M. and S.S. Funding acquisition: S.S.

## Acknowledgements

We thank Elisa Mogollon-Perez, Fabio Dos Santos Barbosa, and Phon Green for their excellent technical assistance. We are also grateful to Matthew Smoker for his expertise on barley transformation and to Alan Wanke for help with initial RNA-seq read alignments. We gratefully acknowledge Mark Youles and TSL SynBio for providing plasmid pICSL90010. We thank all members of the Schornack team for their valuable feedback and discussions throughout the project. We also appreciate Ralph Hückelhoven’s insightful comments and constructive feedback on the data and overall project.

## Funding

This work was funded by UKRI/BBSRC (BB/X014118/1), the Gatsby Charitable Foundation (GAT3395/GLD), the Perry Foundation (G129382) and the European Research Council (ERC-2014-STG, H2020, 637537).

## Conflicts of Interest

The authors declare no competing interests

## Data availability statement

Data deposition: The raw fastq data have been deposited at the European Nucleotide Archive (ENA), https://www.ebi.ac.uk/ena/browser/home (accession no. PRJEB110654). All other data needed to evaluate the conclusions in the paper are present in the paper and/or the Supplementary Materials.

## Notes

### Competing Interest Statement

The authors have declared no competing interest.

## References

Abdelghany, A.M., Lamlom, S.F., and Naser, M. (2024). Dissecting the resilience of barley genotypes under multiple adverse environmental conditions. BMC Plant Biol 24, 16.

Bai, J., Zhu, X., Wang, Q., Zhang, J., Chen, H., Dong, G., Zhu, L., Zheng, H., Xie, Q., Nian, J., Chen, F., Fu, Y., Qian, Q., and Zuo, J. (2015). Rice TUTOU1 Encodes a Suppressor of cAMP Receptor-Like Protein That Is Important for Actin Organization and Panicle Development. Plant Physiol 169, 1179–1191.

Basheer, J., Vadovič, P., Šamajová, O., Melicher, P., Komis, G., Křenek, P., Králová, M., Pechan, T., Ovečka, M., Takáč, T., and Šamaj, J. (2022). Knockout of MITOGEN-ACTIVATED PROTEIN KINASE 3 causes barley root resistance against Fusarium graminearum. Plant Physiol 190, 2847–2867.

Basu, D., Le, J., El-Essal Sel, D., Huang, S., Zhang, C., Mallery, E.L., Koliantz, G., Staiger, C.J., and Szymanski, D.B. (2005). DISTORTED3/SCAR2 is a putative arabidopsis WAVE complex subunit that activates the Arp2/3 complex and is required for epidermal morphogenesis. Plant Cell 17, 502–524.

Berger, F., and Gutjahr, C. (2021). Factors affecting plant responsiveness to arbuscular mycorrhiza. Curr Opin Plant Biol 59, 101994.

Bettgenhaeuser, J., Hernández-Pinzón, I., Dawson, A.M., Gardiner, M., Green, P., Taylor, J., Smoker, M., Ferguson, J.N., Emmrich, P., Hubbard, A., Bayles, R., Waugh, R., Steffenson, B.J., Wulff, B.B.H., Dreiseitl, A., Ward, E.R., and Moscou, M.J. (2021). The barley immune receptor Mla recognizes multiple pathogens and contributes to host range dynamics. Nat Commun 12, 6915.

Brumm, S., Gavrin, A., Macleod, M., Chesneau, G., Uslander, A., and Schornack, S. (2025). Functional divergence of plant SCAR/WAVE proteins is determined by intrinsically disordered regions. Sci Adv 11, eadt6107.

Büschges, R., Hollricher, K., Panstruga, R., Simons, G., Wolter, M., Frijters, A., van Daelen, R., van der Lee, T., Diergaarde, P., Groenendijk, J., Töpsch, S., Vos, P., Salamini, F., and Schulze-Lefert, P. (1997). The barley Mlo gene: a novel control element of plant pathogen resistance. Cell 88, 695–705.

Campo, S., Martín-Cardoso, H., Olivé, M., Pla, E., Catala-Forner, M., Martínez-Eixarch, M., and San Segundo, B. (2020). Effect of Root Colonization by Arbuscular Mycorrhizal Fungi on Growth, Productivity and Blast Resistance in Rice. Rice (N Y) 13, 42.

Capella-Gutierrez, S., Silla-Martinez, J.M., and Gabaldon, T. (2009). trimAl: a tool for automated alignment trimming in large-scale phylogenetic analyses. Bioinformatics 25, 1972–1973.

Carpita, N.C., and Gibeaut, D.M. (1993). Structural models of primary cell walls in flowering plants: consistency of molecular structure with the physical properties of the walls during growth. Plant J 3, 1–30.

Chitarra, W., Pagliarani, C., Maserti, B., Lumini, E., Siciliano, I., Cascone, P., Schubert, A., Gambino, G., Balestrini, R., and Guerrieri, E. (2016). Insights on the Impact of Arbuscular Mycorrhizal Symbiosis on Tomato Tolerance to Water Stress. Plant Physiol 171, 1009–1023.

Choi, J., Summers, W., and Paszkowski, U. (2018). Mechanisms Underlying Establishment of Arbuscular Mycorrhizal Symbioses. Annu Rev Phytopathol 56, 135–160.

Delgado-Baquerizo, M., Guerra, C.A., Cano-Díaz, C., Egidi, E., Wang, J.T., Eisenhauer, N., Singh, B.K., and Maestre, F.T. (2020). The proportion of soil-borne pathogens increases with warming at the global scale. Nature Climate Change 10, 550-+.

Djakovic, S., Dyachok, J., Burke, M., Frank, M.J., and Smith, L.G. (2006). BRICK1/HSPC300 functions with SCAR and the ARP2/3 complex to regulate epidermal cell shape in Arabidopsis. Development 133, 1091–1100.

Dobin, A., Davis, C.A., Schlesinger, F., Drenkow, J., Zaleski, C., Jha, S., Batut, P., Chaisson, M., and Gingeras, T.R. (2013). STAR: ultrafast universal RNA-seq aligner. Bioinformatics 29, 15–21.

Evangelisti, E., Turner, C., McDowell, A., Shenhav, L., Yunusov, T., Gavrin, A., Servante, E.K., Quan, C., and Schornack, S. (2021). Deep learning-based quantification of arbuscular mycorrhizal fungi in plant roots. New Phytol 232, 2207–2219.

Ewels, P.A., Peltzer, A., Fillinger, S., Patel, H., Alneberg, J., Wilm, A., Garcia, M.U., Di Tommaso, P., and Nahnsen, S. (2020). The nf-core framework for community-curated bioinformatics pipelines. Nat Biotechnol 38, 276–278.

FAOSTAT. (2024). Food and Agriculture Organization of the United Nations Statistics Division. Crops and livestock products, World + (Total), production quantity, primary crops, 2024; https://www.fao.org/faostat/en/#data/QCL/visualize data accessed on 09.02.2026.

Fawke, S., Torode, T.A., Gogleva, A., Fich, E.A., Sorensen, I., Yunusov, T., Rose, J.K.C., and Schornack, S. (2019). Glycerol-3-phosphate acyltransferase 6 controls filamentous pathogen interactions and cell wall properties of the tomato and Nicotiana benthamiana leaf epidermis. New Phytol 223, 1547–1559.

Fisher, M.C., Hawkins, N.J., Sanglard, D., and Gurr, S.J. (2018). Worldwide emergence of resistance to antifungal drugs challenges human health and food security. Science 360, 739–742.

Freisleben, R., and Lein, A. (1942). Über die Auffindung einer mehltauresistenten Mutante nach Röntgenbestrahlung einer anfälligen reinen Linie von Sommergerste. Die Naturwissenschaften 30, 608–608.

Garcia-Ruiz, H., Szurek, B., and Van den Ackerveken, G. (2021). Stop helping pathogens: engineering plant susceptibility genes for durable resistance. Curr Opin Biotechnol 70, 187–195.

Gavrin, A., Rey, T., Torode, T.A., Toulotte, J., Chatterjee, A., Kaplan, J.L., Evangelisti, E., Takagi, H., Charoensawan, V., Rengel, D., Journet, E.P., Debelle, F., de Carvalho-Niebel, F., Terauchi, R., Braybrook, S., and Schornack, S. (2020). Developmental Modulation of Root Cell Wall Architecture Confers Resistance to an Oomycete Pathogen. Curr Biol 30, 4165–4176 e4165.

Grace, E.J., Cotsaftis, O., Tester, M., Smith, F.A., and Smith, S.E. (2009). Arbuscular mycorrhizal inhibition of growth in barley cannot be attributed to extent of colonization, fungal phosphorus uptake or effects on expression of plant phosphate transporter genes. New Phytol 181, 938–949.

Gutteridge, R.J., Bateman, G.L., and Todd, A.D. (2003). Variation in the effects of take-all disease on grain yield and quality of winter cereals in field experiments. Pest Manag Sci 59, 215–224.

Guyon, A., Staps, T., Badot, L., and Schornack, S. (2025). Mutualist-pathogen co-colonization modulates phosphoinositide signatures at host intracellular interfaces. Cell Rep 44, 116702.

Hensel, G., Kastner, C., Oleszczuk, S., Riechen, J., and Kumlehn, J. (2009). Agrobacterium-mediated gene transfer to cereal crop plants: current protocols for barley, wheat, triticale, and maize. Int J Plant Genomics 2009, 835608.

Jacott, C.N., Charpentier, M., Murray, J.D., and Ridout, C.J. (2020). Mildew Locus O facilitates colonization by arbuscular mycorrhizal fungi in angiosperms. New Phytol 227, 343–351.

Jakobsen, I., and Rosendahl, L. (1990). Carbon Flow into Soil and External Hyphae from Roots of Mycorrhizal Cucumber Plants. New Phytologist 115, 77–83.

Jarosch, B., Kogel, K.H., and Schaffrath, U. (1999). The ambivalence of the barley locus:: Mutations conferring resistance against powdery mildew (f. sp,) enhance susceptibility to the rice blast fungus. Molecular Plant-Microbe Interactions 12, 508–514.

Jayakodi, M., Lu, Q., Pidon, H., Rabanus-Wallace, M.T., Bayer, M., Lux, T., Guo, Y., Jaegle, B., Badea, A., Bekele, W., Brar, G.S., Braune, K., Bunk, B., Chalmers, K.J., Chapman, B., Jørgensen, M.E., Feng, J.W., Feser, M., Fiebig, A., Gundlach, H., Guo, W., Haberer, G., Hansson, M., Himmelbach, A., Hoffie, I., Hoffie, R.E., Hu, H., Isobe, S., König, P., Kale, S.M., Kamal, N., Keeble-Gagnère, G., Keller, B., Knauft, M., Koppolu, R., Krattinger, S.G., Kumlehn, J., Langridge, P., Li, C., Marone, M.P., Maurer, A., Mayer, K.F.X., Melzer, M., Muehlbauer, G.J., Murozuka, E., Padmarasu, S., Perovic, D., Pillen, K., Pin, P.A., Pozniak, C.J., Ramsay, L., Pedas, P.R., Rutten, T., Sakuma, S., Sato, K., Schüler, D., Schmutzer, T., Scholz, U., Schreiber, M., Shirasawa, K., Simpson, C., Skadhauge, B., Spannagl, M., Steffenson, B.J., Thomsen, H.C., Tibbits, J.F., Nielsen, M.T.S., Trautewig, C., Vequaud, D., Voss, C., Wang, P., Waugh, R., Westcott, S., Rasmussen, M.W., Zhang, R., Zhang, X.Q., Wicker, T., Dockter, C., Mascher, M., and Stein, N. (2024). Structural variation in the pangenome of wild and domesticated barley. Nature 636, 654–662.

Jones, J.D.G., Staskawicz, B.J., and Dangl, J.L. (2024). The plant immune system: From discovery to deployment. Cell 187, 2095–2116.

Jørgensen, J.H. (1992). Discovery, Characterization and Exploitation of Mlo Powdery Mildew Resistance in Barley. Euphytica 63, 141–152.

Jørgensen, J.H., and Wolfe, M. (2011). Genetics of Powdery Mildew Resistance in Barley. Critical Reviews in Plant Sciences 13, 97–119.

Kakouridis, A., Hagen, J.A., Kan, M.P., Mambelli, S., Feldman, L.J., Herman, D.J., Weber, P.K., Pett-Ridge, J., and Firestone, M.K. (2022). Routes to roots: direct evidence of water transport by arbuscular mycorrhizal fungi to host plants. New Phytol 236, 210–221.

Keymer, A., Pimprikar, P., Wewer, V., Huber, C., Brands, M., Bucerius, S.L., Delaux, P.M., Klingl, V., Ropenack-Lahaye, E.V., Wang, T.L., Eisenreich, W., Dormann, P., Parniske, M., and Gutjahr, C. (2017). Lipid transfer from plants to arbuscular mycorrhiza fungi. Elife 6.

Kim, S.J., and Brandizzi, F. (2021). Advances in Cell Wall Matrix Research with a Focus on Mixed-Linkage Glucan. Plant Cell Physiol 62, 1839–1846.

Kjær, B., Jensen, H.P., Jensen, J., and Jørgensen, J.H. (1990). Associations between three ml-o powdery mildew resistance genes and agronomic traits in barley. Euphytica 46, 185–193.

Kollmar, M., Lbik, D., and Enge, S. (2012). Evolution of the eukaryotic ARP2/3 activators of the WASP family: WASP, WAVE, WASH, and WHAMM, and the proposed new family members WAWH and WAML. BMC Res Notes 5, 88.

Kurihara, D., Mizuta, Y., Sato, Y., and Higashiyama, T. (2015). ClearSee: a rapid optical clearing reagent for whole-plant fluorescence imaging. Development 142, 4168–4179.

Lazo, G.R., Stein, P.A., and Ludwig, R.A. (1991). A DNA transformation-competent Arabidopsis genomic library in Agrobacterium. Biotechnology (N Y) 9, 963–967.

Le Fevre, R., O’Boyle, B., Moscou, M.J., and Schornack, S. (2016). Colonization of Barley by the Broad-Host Hemibiotrophic Pathogen Phytophthora palmivora Uncovers a Leaf Development-Dependent Involvement of Mlo. Mol Plant Microbe Interact 29, 385–395.

Letunic, I., and Bork, P. (2024). Interactive Tree of Life (iTOL) v6: recent updates to the phylogenetic tree display and annotation tool. Nucleic Acids Res 52, W78–W82.

Li, Y., Sorefan, K., Hemmann, G., and Bevan, M.W. (2004). Arabidopsis NAP and PIR regulate actin-based cell morphogenesis and multiple developmental processes. Plant Physiol 136, 3616–3627.

Limpens, E., Mirabella, R., Fedorova, E., Franken, C., Franssen, H., Bisseling, T., and Geurts, R. (2005). Formation of organelle-like N2-fixing symbiosomes in legume root nodules is controlled by DMI2. Proc Natl Acad Sci U S A 102, 10375–10380.

Liu, Y., Esposto, D., Mahdi, L.K., Porzel, A., Stark, P., Hussain, H., Scherr-Henning, A., Isfort, S., Bathe, U., Acosta, I.F., Zuccaro, A., Balcke, G.U., and Tissier, A. (2024). Hordedane diterpenoid phytoalexins restrict Fusarium graminearum infection but enhance Bipolaris sorokiniana colonization of barley roots. Mol Plant 17, 1307–1327.

Love, M.I., Huber, W., and Anders, S. (2014). Moderated estimation of fold change and dispersion for RNA-seq data with DESeq2. Genome Biol 15, 550.

Luginbuehl, L.H., Menard, G.N., Kurup, S., Van Erp, H., Radhakrishnan, G.V., Breakspear, A., Oldroyd, G.E.D., and Eastmond, P.J. (2017). Fatty acids in arbuscular mycorrhizal fungi are synthesized by the host plant. Science 356, 1175–1178.

Macleod, M., Brumm, S., and Schornack, S. (2025). Quantification of Phytophthora palmivora Infection in Barley and Related Monocot Roots. Methods Mol Biol 2892, 105–116.

McDonald, B.A., and Linde, C. (2002). Pathogen population genetics, evolutionary potential, and durable resistance. Annu Rev Phytopathol 40, 349–379.

Miyahara, A., Richens, J., Starker, C., Morieri, G., Smith, L., Long, S., Downie, J.A., and Oldroyd, G.E. (2010). Conservation in function of a SCAR/WAVE component during infection thread and root hair growth in Medicago truncatula. Mol Plant Microbe Interact 23, 1553–1562.

Newton, A.C., Flavell, A.J., George, T.S., Leat, P., Mullholland, B., Ramsay, L., Revoredo-Giha, C., Russell, J., Steffenson, B.J., Swanston, J.S., Thomas, W.T.B., Waugh, R., White, P.J., and Bingham, I.J. (2011). Crops that feed the world 4. Barley: a resilient crop? Strengths and weaknesses in the context of food security. Food Security 3, 141–178.

Nguyen, L.T., Schmidt, H.A., von Haeseler, A., and Minh, B.Q. (2015). IQ-TREE: a fast and effective stochastic algorithm for estimating maximum-likelihood phylogenies. Mol Biol Evol 32, 268–274.

Pathuri, I.P., Zellerhoff, N., Schaffrath, U., Hensel, G., Kumlehn, J., Kogel, K.H., Eichmann, R., and Huckelhoven, R. (2008). Constitutively activated barley ROPs modulate epidermal cell size, defense reactions and interactions with fungal leaf pathogens. Plant Cell Rep 27, 1877–1887.

Patro, R., Duggal, G., Love, M.I., Irizarry, R.A., and Kingsford, C. (2017). Salmon provides fast and bias-aware quantification of transcript expression. Nat Methods 14, 417–419.

Rey, T., Chatterjee, A., Buttay, M., Toulotte, J., and Schornack, S. (2015). Medicago truncatula symbiosis mutants affected in the interaction with a biotrophic root pathogen. New Phytol 206, 497–500.

Rozewicki, J., Li, S., Amada, K.M., Standley, D.M., and Katoh, K. (2019). MAFFT-DASH: integrated protein sequence and structural alignment. Nucleic Acids Res 47, W5–W10.

Sawers, R.J., Svane, S.F., Quan, C., Gronlund, M., Wozniak, B., Gebreselassie, M.N., Gonzalez-Munoz, E., Chavez Montes, R.A., Baxter, I., Goudet, J., Jakobsen, I., and Paszkowski, U. (2017). Phosphorus acquisition efficiency in arbuscular mycorrhizal maize is correlated with the abundance of root-external hyphae and the accumulation of transcripts encoding PHT1 phosphate transporters. New Phytol 214, 632–643.

Scheler, B., Schnepf, V., Galgenmuller, C., Ranf, S., and Huckelhoven, R. (2016). Barley disease susceptibility factor RACB acts in epidermal cell polarity and positioning of the nucleus. J Exp Bot 67, 3263–3275.

Schroeder, K.L., and Paulitz, T.C. (2006). Root Diseases of Wheat and Barley During the Transition from Conventional Tillage to Direct Seeding. Plant Dis 90, 1247–1253.

Teillet, A., Garcia, J., de Billy, F., Gherardi, M., Huguet, T., Barker, D.G., de Carvalho-Niebel, F., and Journet, E.P. (2008). api, A novel Medicago truncatula symbiotic mutant impaired in nodule primordium invasion. Mol Plant Microbe Interact 21, 535–546.

Thirkell, T.J., Campbell, M., Driver, J., Pastok, D., Merry, B., and Field, K.J. (2021). Cultivar-dependent increases in mycorrhizal nutrient acquisition by barley in response to elevated CO. Plants People Planet 3, 553–566.

Tkacz, A., Ledermann, R., Martyn, A., Schornack, S., Oldroyd, G.E.D., and Poole, P.S. (2022). Nodulation and nitrogen fixation in Medicago truncatula strongly alters the abundance of its root microbiota and subtly affects its structure. Environ Microbiol 24, 5524–5533.

Uhrig, J.F., Mutondo, M., Zimmermann, I., Deeks, M.J., Machesky, L.M., Thomas, P., Uhrig, S., Rambke, C., Hussey, P.J., and Hülskamp, M. (2007). The role of Arabidopsis SCAR genes in ARP2-ARP3-dependent cell morphogenesis. Development 134, 967–977.

van Schie, C.C., and Takken, F.L. (2014). Susceptibility genes 101: how to be a good host. Annu Rev Phytopathol 52, 551–581.

Veltman, D.M., and Insall, R.H. (2010). WASP family proteins: their evolution and its physiological implications. Mol Biol Cell 21, 2880–2893.

Vierheilig, H., Coughlan, A.P., Wyss, U., and Piche, Y. (1998). Ink and vinegar, a simple staining technique for arbuscular-mycorrhizal fungi. Appl Environ Microbiol 64, 5004–5007.

Yanagisawa, M., Zhang, C., and Szymanski, D.B. (2013). ARP2/3-dependent growth in the plant kingdom: SCARs for life. Front Plant Sci 4, 166.

Zhang, C., Mallery, E.L., Schlueter, J., Huang, S., Fan, Y., Brankle, S., Staiger, C.J., and Szymanski, D.B. (2008). Arabidopsis SCARs function interchangeably to meet actin-related protein 2/3 activation thresholds during morphogenesis. Plant Cell 20, 995–1011.

