## Supplementary Figures and Tables for "Engineering quantitative root disease resistance in barley by targeting conserved SCAR susceptibility genes without compromising seed yield or mycorrhizal symbiosis"

Sabine Brumm et al.

*Sabine Brumm. and Sebastian Schornack.

This PDF file includes:
Figure S1 to S7
Table S1 and S2

Other Supplementary Materials for this manuscript include the following:
Data S1 to S3

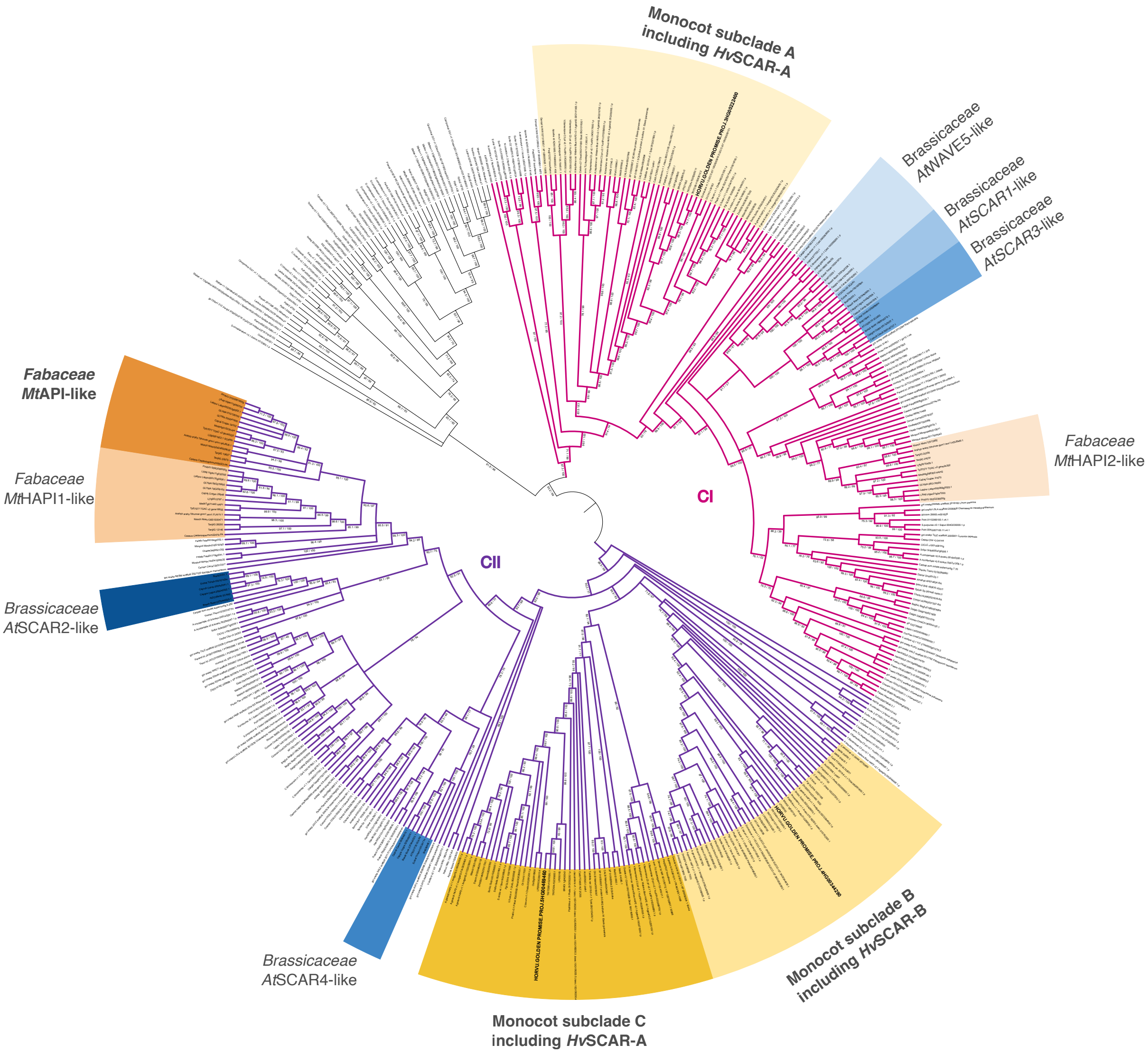


**Figure S1: SCAR/WAVE proteins are conserved across the plant kingdom**
Maximum-likelihood phylogeny of plant SCAR/WAVE proteins. Angiosperm SCAR/WAVE proteins cluster into two major clades, CI (pink) and CII (lilac). Within these clades, lineage-specific subclades are highlighted. In Brassicaceae, *At*WAVE5-, *At*SCAR1-, *At*SCAR3-, *At*SCAR4-, and *At*SCAR2-like subclades are shown in shades of blue. In Fabaceae, *Mt*API-, *Mt*HAPI1-, and *Mt*HAPI2-like subclades are highlighted in shades of orange. In monocots, three distinct subclades (A, B, and C) were identified and are highlighted in shades of yellow. Each monocot subclade contains one barley gene, with barley identifiers shown in bold.

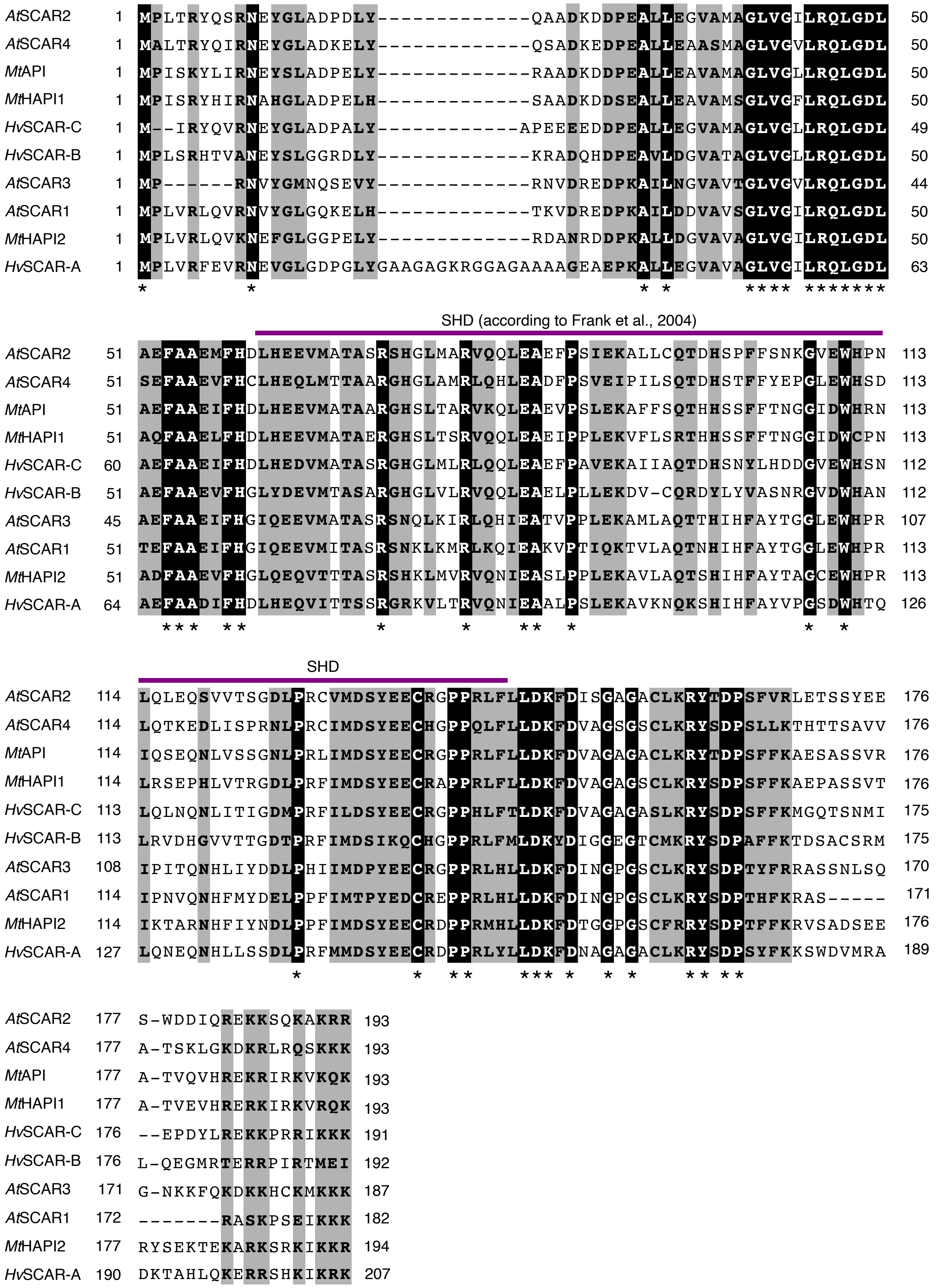


**Figure S2: MAFFT alignment of the SCAR SHD domains from *A. thaliana*, *M. truncatula*, and barley.** Sequences were aligned using the EMBL MAFFT algorithm, and similarity information was obtained with ESPript3 based on the BLOSUM62 scoring matrix (global score threshold: 0.2, no grouping). Identical residues are highlighted with black boxes, white bold characters, and an asterisk below the alignment. Similar residues are indicated by bold characters within grey boxes. The SHD (SCAR homology domains, purple bar) are annotated according to (Frank et al., 2004).

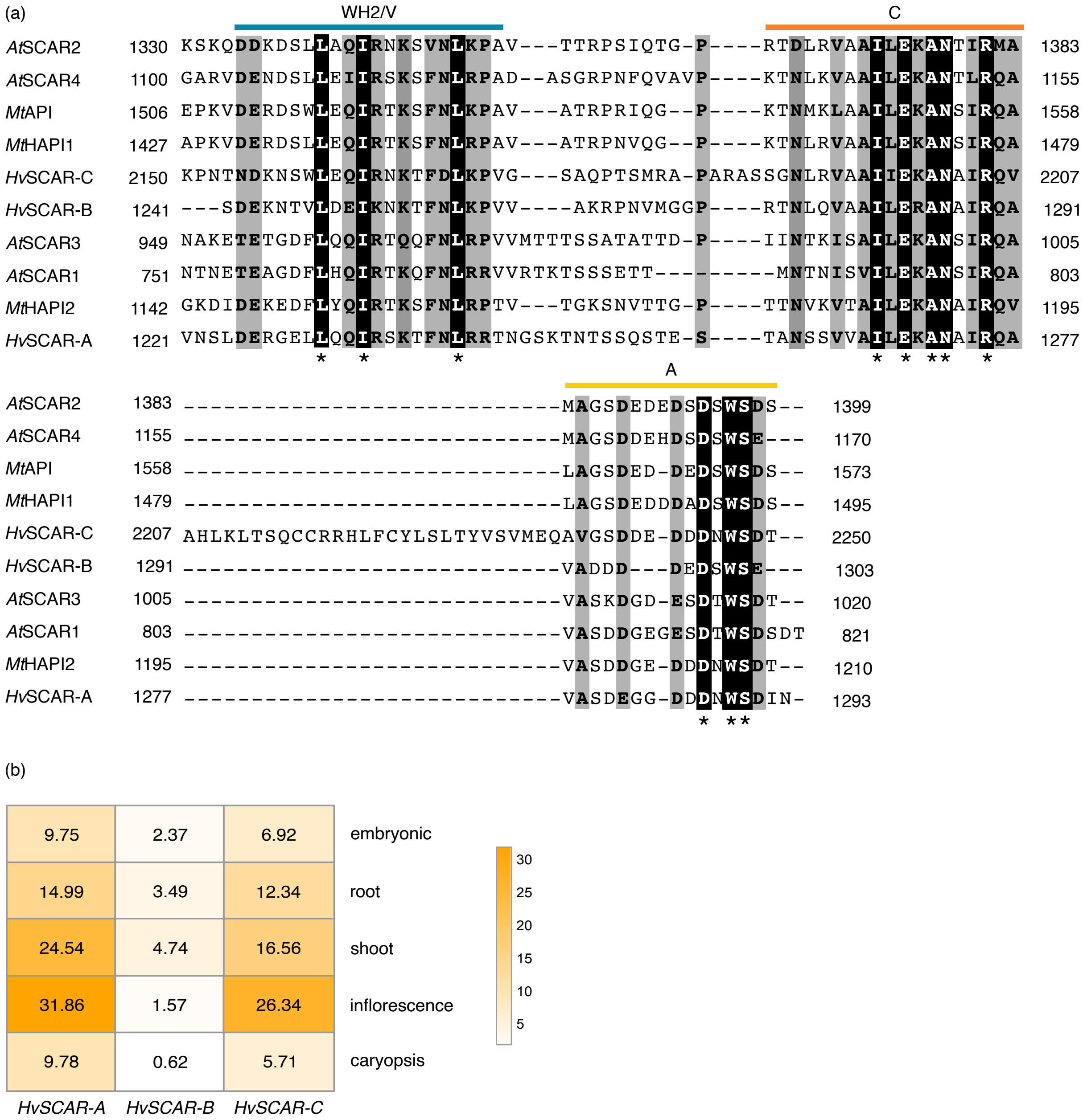


**Figure S3: MAFFT alignment of the SCAR WA domains and *HvSCAR* expression data.** a) MAFFT alignment of the SCAR WCA (WASP homology 2, central, acidic) domains from *A. thaliana*, *M. truncatula*, and barley**.** Sequences were aligned using the EMBL MAFFT algorithm, and similarity information was obtained with ESPript3 based on the BLOSUM62 scoring matrix (global score threshold: 0.2, no grouping). Identical residues are highlighted with black boxes, white bold characters, and an asterisk below the alignment. Similar residues are indicated by bold characters within grey boxes. The WH2/V (WASP homology 2/Verprolin) (blue bar), C (orange bar) and A (Acidic) domain (yellow bar) are annotated according to (Frank et al., 2004). b) Heatmap representation of expression levels as transcripts per million obtained for *HvSCAR-A*, *HvSCAR-B* and *HvSCAR-C* from Panbarlex for the accession Golden Promise.

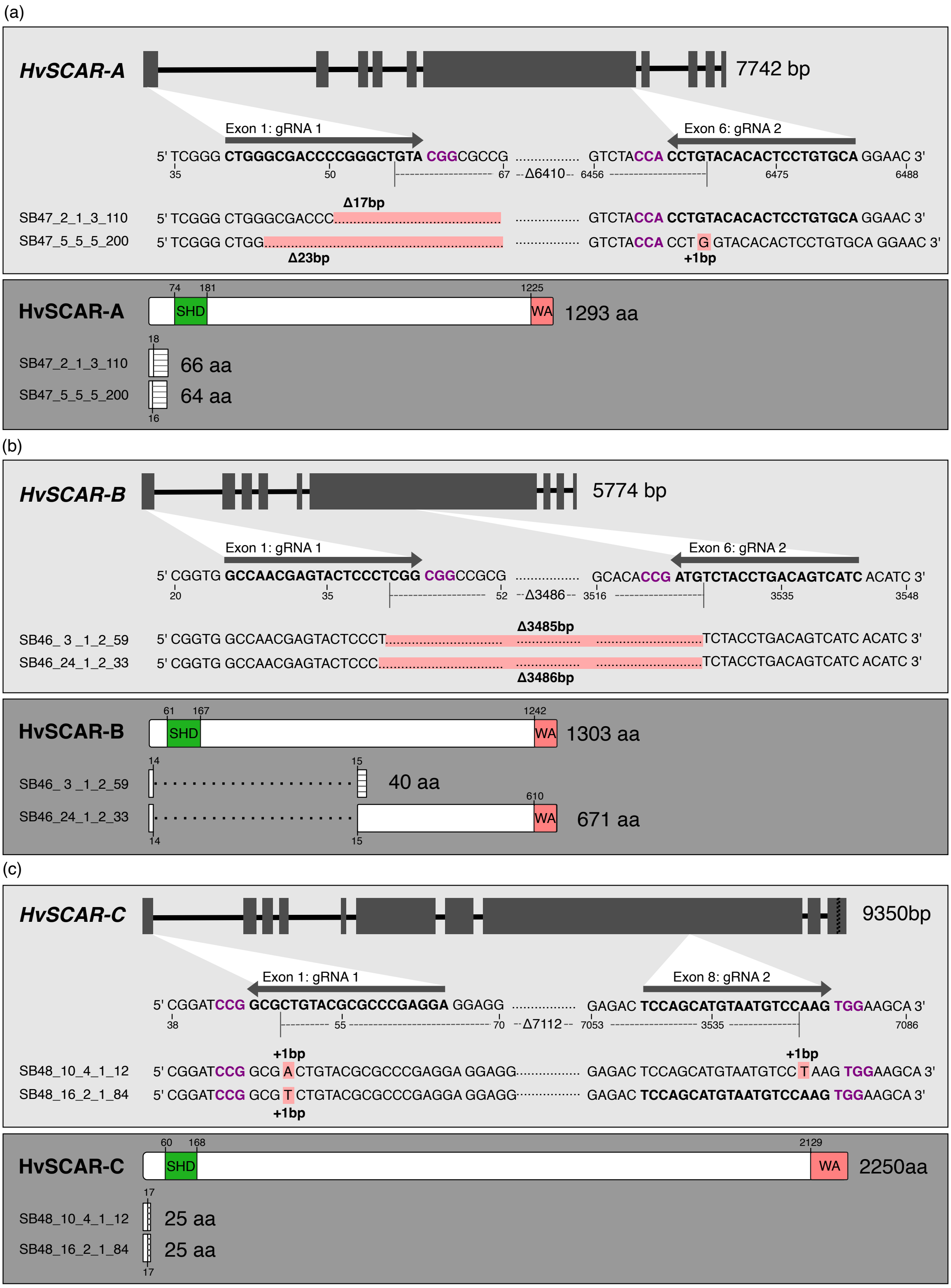


**Figure S4. CRISPR-induced mutations in *HvSCAR* genes and their predicted impact on protein translation.** Panels show schematic representation of mutations in two independent knockout lines for *HvSCAR-A* (a), *HvSCAR-B* (b) and *HvSCAR-C* (c). Top of each panel (light grey boxes): Gene structure schematics showing exon–intron organization, guide RNA positions (white triangles), and detailed gRNA sequences with corresponding genomic modifications. Positional information is given relative to the ATG start codon of each gene. Bottom of each panel (dark grey boxes): Schematic representation of SCAR proteins, highlighting conserved domains: N-terminal SHD (green) and C-terminal WA domain (red). Amino acid positions are indicated. Changes in amino acid sequences are shown as lines within white boxes, and the predicted length of the resulting peptides is provided.

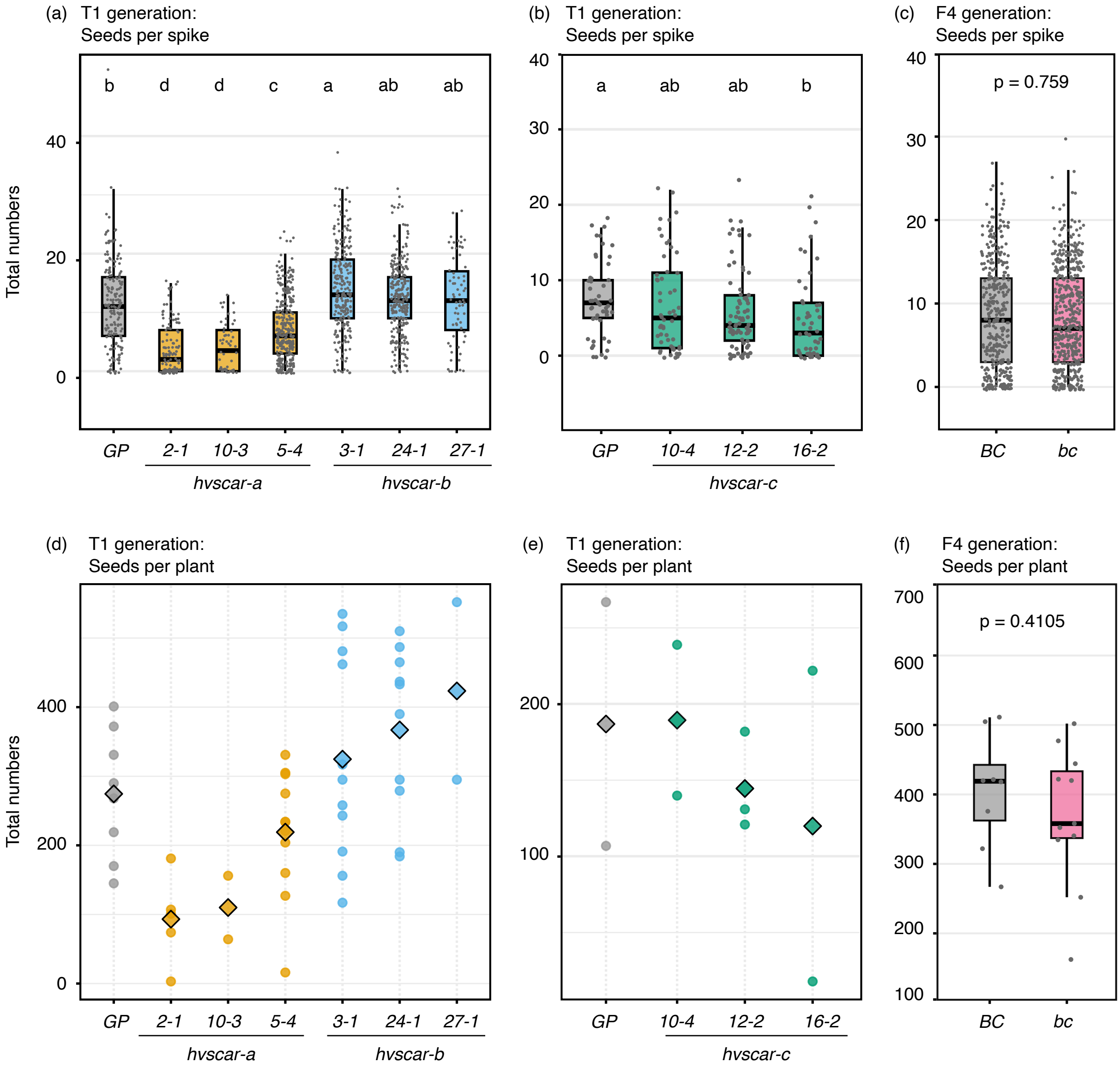


**Figure S5: Seed production across different plant generations.** Harvest data from T1 generation *hvscar* single CRISPR knockout mutants and F4 generation *hvscar-b,c* double mutants are shown. Within each experiment, mutants and their respective controls were grown under identical conditions. However, the three datasets originate from different time points and growth chambers. (a–c) Boxplot representation of number of seeds per spike. Each data point represents the number of seeds harvested from a single spike. (a) T1 *hvscar-a* lines *2-1* (n = 131), *10-3* (n = 54), and *5-4* (n = 342), and T1 *hvscar-b* lines *3-1* (n = 254), *24-1* (n = 296), and *27-1* (n = 69) were compared to Golden Promise (n = 191). (b) T1 *hvscar-c* lines *10-4* (n = 77), *12-2* (n = 55), and *16-2* (n = 49) were compared to Golden Promise (n = 50). (c) F4 *hvscar-b,c* double mutant (n = 495) was compared to the *HvSCAR-B,C* control (n = 383). (d–f) Dotplot (d,e) and boxplot (f) representation of number of seeds per plant. Each data point represents the total number of seeds per individual plant. Diamonds in the dotplots represent the mean. (d) T1 *hvscar-a* lines *2-1* (n = 5), *10-3* (n = 2), and *5-4* (n = 11), and T1 *hvscar-b* lines *3-1* (n = 11), *24-1* (n = 10), and *27-1* (n = 2) were compared to Golden Promise (n = 8). (e) T1 *hvscar-c* lines *10-4* (n = 2), *12-2* (n = 3), and *16-2* (n = 2) were compared to Golden Promise (n = 2). (f) F4 *hvscar-b,c* double mutants (n = 11) were compared to the *HvSCAR-B,C* control (n = 8). Statistical analysis: For datasets with ≥3 data points per group, normality was assessed using the Shapiro-Wilk test. Depending on data distribution and variance, statistical differences were determined using Kruskal-Wallis tests followed by Bonferroni correction (a, b), Wilcoxon rank-sum test (c), or Welch's t-test (f). Significant differences are indicated by different letters for multiple comparisons or exact p-values are shown within the respective panels.

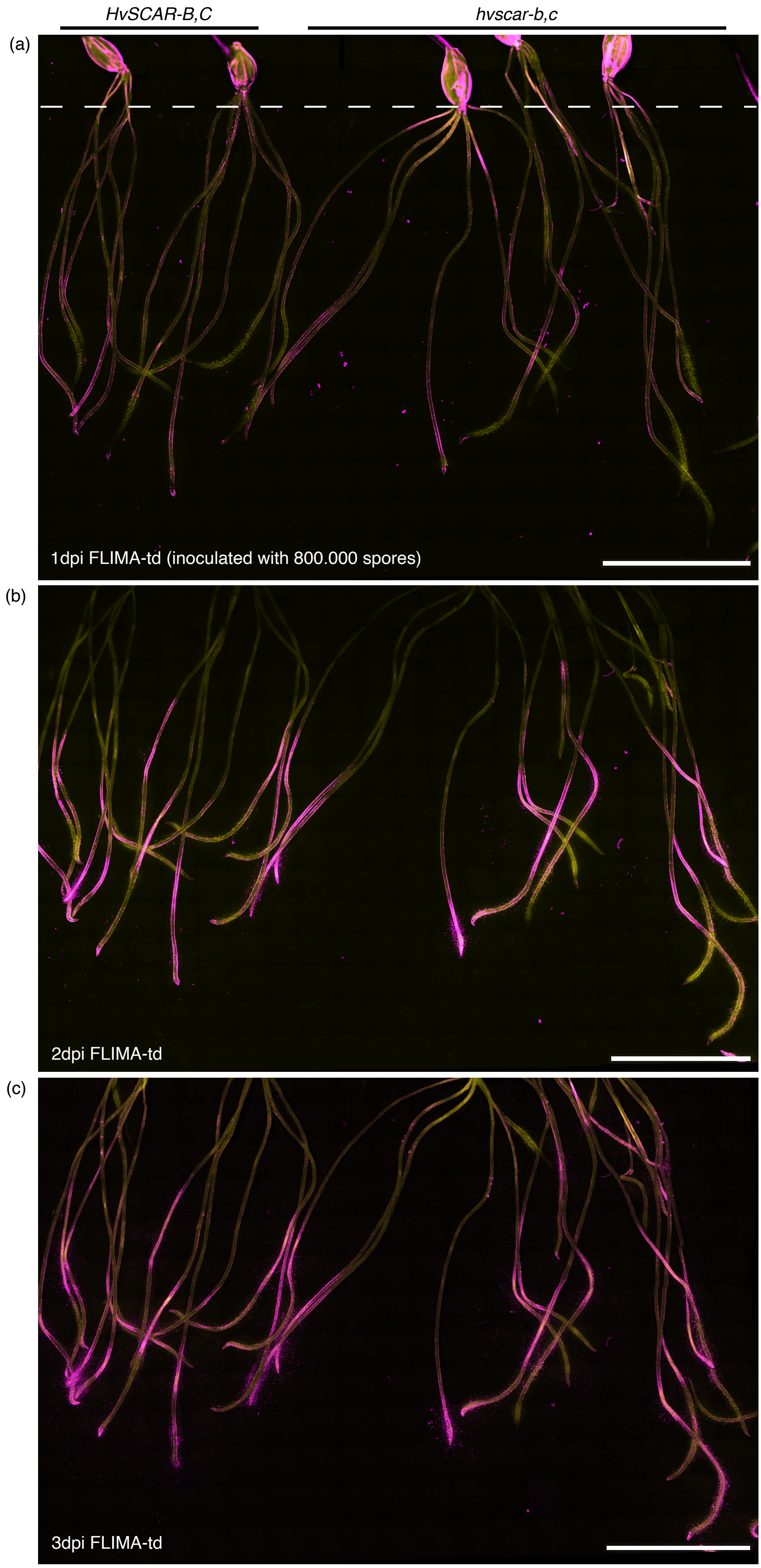
HvSCAR-B,C
hvscar-b,c
(a)
1dpi FLIMA-td (inoculated with 800.000 spores)
(b)
2dpi FLIMA-td
(c)
3dpi FLIMA-td

**Figure S6: Time-course of FLIMA-td infection on hydroponically grown barley roots** (a-b) Representative images of *HvSCAR-B,C* (wild type) and *hvscar-b,c* roots one (a), two (c) and three (c) days after inoculation with *Phytophthora palmivora* strain FLIMA-td. Two *HvSCAR-B,C* and three *hvscar-b,c* seedlings were inoculated with 40.000 spores/ml in 20ml medium (inoculation with 800.000 spores in total). Magenta: FLIMA-td fluorescence. Yellow: autofluorescence barley roots. White dahed line: image clipping in (b) and (c). Scale bar: 20 mm.

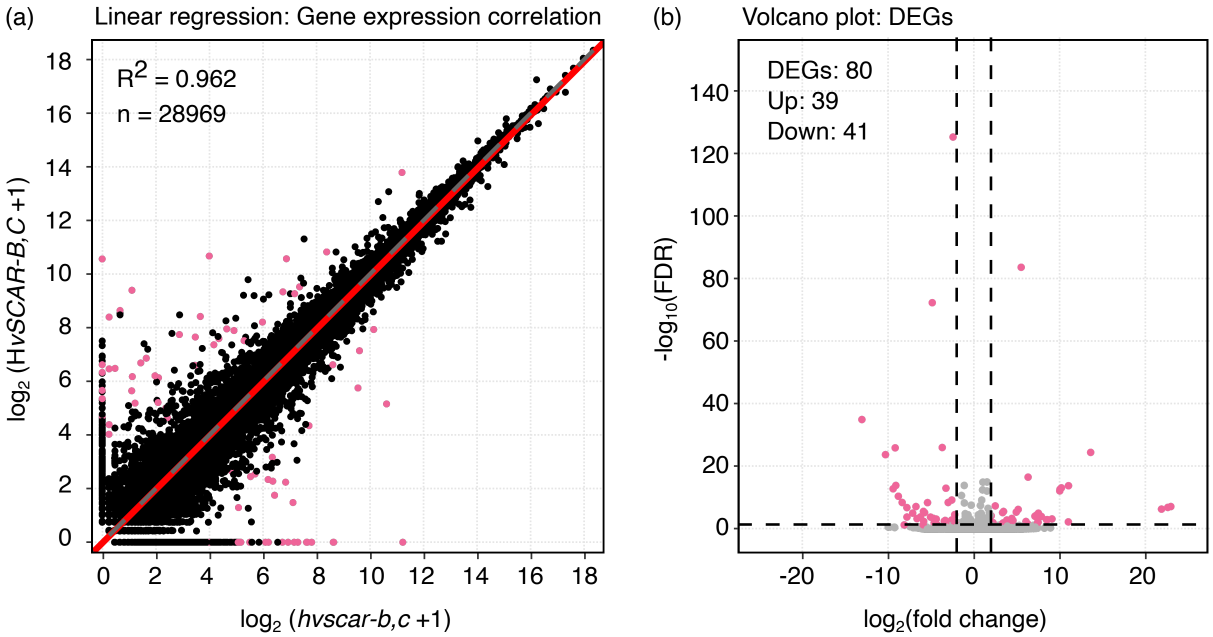


**Supplementary Figure 7: Differential expression analysis of non-infected barley roots**
(a) Low-abundance genes were removed by excluding all genes with a total count ≤ 1 across all samples. For the remaining genes (n = 28,969), a global expression correlation analysis was performed using $log_2$-transformed (pseudocount = 1) DESeq2-normalized mean counts from uninfected *HvSCAR-B,C* and *hvscar-b,c* samples. A linear regression model (*y* ~ *x*) was fitted to the data, and the coefficient of determination ($R^2$) was calculated to quantify the strength of the expression correlation. The red line represents the fitted linear regression between *HvSCAR-B,C* and *hvscar-b,c* expression, while the grey dashed line indicates the identity line (*y* = *x*). Pink dots**:** significantly differentially expressed genes (padj < 0.05 & $|log_2FC| > 2$). Black dots: non-significant genes. (b) Differential gene expression analysis was performed using DESeq2 after filtering out low-abundance genes (total count ≤ 1 across all samples). The volcano plot displays all genes as a function of $log_2$ fold change (x-axis) and $-log_{10}$ adjusted *p*-value (FDR; y-axis). Vertical dashed lines indicate the fold-change threshold ($|log_2FC| = 2$), and the horizontal dashed line marks the significance cutoff (padj = 0.05). Pink dots represent significantly differentially expressed genes (DEGs, padj < 0.05 and $|log_2FC| \geq 2$). Grey dots represent non-significant genes. The numbers of total DEGs, upregulated, and downregulated transcripts are shown in the upper left corner of the plot (also see Data S3).

### Table S1: Primer sequences mentioned in this study

| ID | Name | Primer sequence | Source |
|---|---|---|---|
| SB101 | GatewayFL-HORVU0Hr1G014100.2-F | GGGGACAAGTTTGTACaaaaaagcaggct ATGATACGGTACCAGGTGCGGA | This study |
| SB102 | GatewayFL-HORVU0Hr1G014100.2-stop-R | GGGGACCACTTTGTAcaagaaagctgggtt CTATGTGTCGCTCCAATTATCGTC | This study |
| SB103 | GatewayFL-HORVU3HR1G029210-F | GGGGACAAGTTTGTACaaaaaagcaggct ATGCCGCTGGTCAGGTTCGAGG | This study |
| SB104 | GatewayFL-HORVU3HR1G029210-stop-R | GGGGACCACTTTGTAcaagaaagctgggtt TCAGTTGATATCGCTCCAGTTATC | This study |
| SB30 | PpEF1α-F | CAAGATCCCGTTCGTGCCTA | (Le Fevre et al., 2016) |
| SB31 | PpEF1α-R | GCGTTCAGGTTGTCAAGAGC | (Le Fevre et al., 2016) |
| SB32 | HvEF1a-F | ATGATTCCCACCAAGCCCAT | (Le Fevre et al., 2016) |
| SB33 | HvEF1α-R | ACACCAACAGCCACAGTTTGC | (Le Fevre et al., 2016) |
| SB37 | PpWS21-F | CTCCAGAACGTGTACATTCG | (Rey et al., 2017) |
| SB38 | PpWS21-R | TGGCACCCTTCTCCTCGG | (Rey et al., 2017) |
| SB41 | HvCyc-F | TTGAGGACGAGATAAGGCCAG | |
| SB42 | HvCyc-R | GCGACTGACAAGGTGCAAGAG | |
| SB284 | PpCdc14-F | TCTGCACGAGTTCCAGCATT | (Le Fevre et al., 2016) |
| SB285 | PpCdc14-R | CACCACTAGCGTCACGTTCT | (Le Fevre et al., 2016) |
| SB88 | sgRNA1-HORVU4Hr1G055030-F | aaggtctcaCTTGCCAACGAGTACTCCC TCGGgtttaagagctatgctggaaacag | This study |
| SB90 | sgRNAsBarley-R | tgtggtctctAGCGaaaaaaagcaccgactcggtgccac | This study |
| SB93 | sgRNA1-3Hr1G029210-F | aaggtctcaCTTGCTGGGCGACCCCG GGCTGTAgtttaagagctatgctggaaacag | This study |

| | | | |
|---|---|---|---|
| SB99 | sgRNA1-HORV0Hr1g0141000-F | aaggtctcaCTTGTCCTCGGGCGCGT ACAGCGCgtttaagagctatgctggaaacag | This study |
| SB100 | sgRNA2-HORV0Hr1g0141000-F | aaggtctcaCTTGTCCAGCATGTAATG TCCAAGgtttaagagctatgctggaaacag | This study |
| SB107 | sgRNA2NEW-HORVU3Hr1G029210-F | aaggtctcaCTTGTGCACAGGAGTGTG TACAGGgtttaagagctatgctggaaacag | This study |
| SB108 | sgRNA2AccI-HORVU4HR1G055030-F | aaggtctcaCTTGGATGACTGTCAGGTA GACATgtttaagagctatgctggaaacag | This study |
| SB207 | 3Hr029210CG2_R | CTGGGAAAGTAGCCGTTG | This study |
| SB162 | Hv1g029210_s8R | ACACCTCAACTCCAGTAAAC | This study |
| SB160 | Hv1g029210_s6F | AGGACAATGAGACAAACAAC | This study |
| SB184 | Hv029210CG-R | GAAAGTAGCCAGAGTGAGAG | This study |
| SB180 | Hv055030CGF | AATCAACAGAGAAAAGCTGC | This study |
| SB85 | HORVU4Hr1G055030-Seq4/SB | AAGATGATGTTGGTTCAAGG | This study |
| SB181 | Hv055030CGR | TGTGAGAGTGAATGGAGTTAC | This study |
| SB185 | Hv0141000CG-F | CTGCGAGTGGAGTAAAGC | This study |
| SB186 | Hv0141000CGhae-R | AACCACCCTCCCAGAATAC | This study |
| SB209 | 0Hr014100CG2_R | CACCTGGTACATCAATCTTATC | This study |
| SB187 | HV0141000CGLpn-F | TTGCTCCATGCCCCTTAA | This study |
| SB233 | Cas9_f1 | ATCAATGGGATCCGAGACAA | This study |
| SB234 | Cas9_r1 | AGCCGATTGATGTCCAGTTC | This study |

### Table S2: CDS and vector constructs mentioned in this study

| Name | Description | Source |
|---|---|---|
| pKGW_RR_MGW | Whole vector sequence of multisite Gateway-compatible destination vector with DsRed casette | (Gavrin et al., 2020) |
| pENTR4_1_prMtAPI | Whole vector sequence of pENTR vector with 2kb *M.truncatula API* promoter flanked by attL4 and attR1 sites | (Gavrin et al., 2020) |
| pENTR_p2rp3_T35STerm | Whole vector sequence of pENTR vector with T35S terminator flanked by attR2 and attL3 sites | (Gavrin et al., 2020) |
| HORVU.GOLDEN_PROMISE.PROJ.3HG00222400_+3bp_HvSCAR-A_CDS | Coding sequence of barley HvSCAR-A (HORVU.GOLDEN_PROMISE. PROJ.3HG00222400), modified to include CAT (720-722) as amplified from GP roots (3885bp) | This paper |
| HORVU.GOLDEN_PROMISE.PROJ.4HG00344290_HvSCAR-B_CDS | Coding sequence of barley HvSCAR-B (HORVU.GOLDEN_PROMISE. PROJ.4HG00344290) (3912bp) | This paper |
| HORVU.GOLDEN_PROMISE.PROJ.5HG00469460_-87bp_ HvSCAR-C_CDS | Coding sequence of barley HvSCAR-C (HORVU.GOLDEN_PROMISE. PROJ.5HG00469460), modified to exclude 87bp (in HORVU.GOLDEN_PROMISE .PROJ.5HG00469460 CDS position 6620 to 6706) as amplified from GP roots (6666bp) | This paper |
| SB88_pDONR_HvSCARA | Whole vector sequence of pDONR221 vector with 3885pb HvSCAR-A CDS | This paper |
| SB98_pUC57(Amp)_HvSCAR-B | Whole vector sequence of pUC57 vector with 3912pb HvSCAR-B CDS flanked by attL1 and attL2 sites | This paper |
| SB95_pDONR_HvSCARC | Whole vector sequence of pDONR221 | This paper |

| | | |
|---|---|---|
| | vector with 6666bp HvSCAR-C CDS | |
| SB118_pKGW_pMtAPI_CDS_ HvSCAR-A_35STerm | Binary vector harboring pAPI:: HvSCAR-A::35ST, a pAtUBQ10::DsRed::nos and pnos::KAN::nos expression cassettes | This paper |
| SB115_pKGW_pMtAPI_CDS_ HvSCAR-B_35STerm | Binary vector harboring pAPI:: HvSCAR-B::35ST, a pAtUBQ10::DsRed::nos and pnos::KAN::nos expression cassettes | This paper |
| SB104_pKGW_pMtAPI_CDS_ HvSCARC_35STerm | Binary vector harboring pAPI:: HvSCAR-C::35ST, a pAtUBQ10::DsRed::nos and pnos::KAN::nos expression cassettes | This paper |
| pICSL90010 | gRNA scaffold template plasmid | Unpublish ed |
| pICSL90003 | Level 0 Golden Gate Part: Promoter U6 (Triticum aestivum) | (Lawrens on et al., 2015) Addgene #68262 |
| pICH47751 | Level 1 position-3 | (Weber et al., 2011) Addgene #48002 |
| pICH47761 | Level 1 position-4 | (Weber et al., 2011) Addgene #48003 |
| pICH50900 | Level-M end-linker | (Weber et al., 2011) Addgene #48047 |
| pICSL11059 | Level 1 position 1 35S::hptII::35S hygromycin resistance cassette | (Lawrens on et al., 2015) Addgene #68263 |
| pICSL11056 | Level 1 position 2 Zm::Cas9::nos cassette | (Lawrens on et al., 2015) Addgene #68258 |
| pAGM8031 | Level-M backbone | (Weber et al., 2011) Addgene #48037 |
| SB47_pAGM8031_hvscar_a_ sgRNAs | Binary CRISPR/Cas9 plant transformation vector harboring HvSCAR-A–specific guide RNAs, a Cas9 nuclease cassette, and a hygromycin resistance | This paper |

| | | |
|---|---|---|
| | (HYG) marker for selection. | |
| SB46_pAGM8031_hvscar_b_ sgRNAs | Binary CRISPR/Cas9 plant transformation vector harboring HvSCAR-B–specific guide RNAs, a Cas9 nuclease cassette, and a hygromycin resistance (HYG) marker for selection. | This paper |
| SB48_pAGM8031_hvscar_c_ sgRNAs | Binary CRISPR/Cas9 plant transformation vector harboring HvSCAR-C–specific guide RNAs, a Cas9 nuclease cassette, and a hygromycin resistance (HYG) marker for selection. | This paper |

### Supporting references


**Frank, M., Egile, C., Dyachok, J., Djakovic, S., Nolasco, M., Li, R., and Smith, L.G.** (2004). Activation of Arp2/3 complex-dependent actin polymerization by plant proteins distantly related to Scar/WAVE. Proc Natl Acad Sci U S A **101,** 16379–16384.

**Gavrin, A., Rey, T., Torode, T.A., Toulotte, J., Chatterjee, A., Kaplan, J.L., Evangelisti, E., Takagi, H., Charoensawan, V., Rengel, D., Journet, E.P., Debelle, F., de Carvalho-Niebel, F., Terauchi, R., Braybrook, S., and Schornack, S.** (2020). Developmental Modulation of Root Cell Wall Architecture Confers Resistance to an Oomycete Pathogen. Curr Biol **30,** 4165–4176 e4165.

**Lawrenson, T., Shorinola, O., Stacey, N., Li, C., Østergaard, L., Patron, N., Uauy, C., and Harwood, W.** (2015). Induction of targeted, heritable mutations in barley and Brassica oleracea using RNA-guided Cas9 nuclease. Genome Biol **16,** 258.

**Le Fevre, R., O'Boyle, B., Moscou, M.J., and Schornack, S.** (2016). Colonization of Barley by the Broad-Host Hemibiotrophic Pathogen Phytophthora palmivora Uncovers a Leaf Development-Dependent Involvement of Mlo. Mol Plant Microbe Interact **29,** 385–395.

**Rey, T., Bonhomme, M., Chatterjee, A., Gavrin, A., Toulotte, J., Yang, W., André, O., Jacquet, C., and Schornack, S.** (2017). The Medicago truncatula GRAS protein RAD1 supports arbuscular mycorrhiza symbiosis and Phytophthora palmivora susceptibility. J Exp Bot **68,** 5871–5881.

**Weber, E., Engler, C., Gruetzner, R., Werner, S., and Marillonnet, S.** (2011). A modular cloning system for standardized assembly of multigene constructs. PLoS One **6,** e16765.
